# Synergistic Binding of the Halide and Cationic Prime Substrate of the l-Lysine 4-Chlorinase, BesD, in Both Ferrous and Ferryl States

**DOI:** 10.1101/2023.05.02.539147

**Authors:** Jeffrey W. Slater, Monica E. Neugebauer, Molly J. McBride, Debangsu Sil, Chi-Yun Lin, Bryce J. Katch, Amie K. Boal, Michelle C.Y. Chang, Alexey Silakov, Carsten Krebs, J. Martin Bollinger

## Abstract

An aliphatic halogenase requires four substrates: 2-oxoglutarate (2OG), halide (Cl^−^ or Br^−^), the halogenation target (“prime substrate”), and dioxygen. In well-studied cases, the three non-gaseous substrates must bind to activate the enzyme’s Fe(II) cofactor for efficient capture of O_2_. Halide, 2OG, and (lastly) O_2_ all coordinate directly to the cofactor to initiate its conversion to a *cis*-halo-oxo-iron(IV) (haloferryl) complex, which abstracts hydrogen (H•) from the non-coordinating prime substrate to enable radicaloid carbon-halogen coupling. We dissected the kinetic pathway and thermodynamic linkage in binding of the first three substrates of the l-lysine 4-chlorinase, BesD. After 2OG adds, subsequent coordination of the halide to the cofactor and binding of cationic l-Lys near the cofactor are associated with strong heterotropic cooperativity. Progression to the haloferryl intermediate upon addition of O_2_ does not trap the substrates in the active site and, in fact, markedly diminishes cooperativity between halide and l-Lys. The surprising lability of the BesD•[Fe(IV)=O]•Cl•succinate•l-Lys complex engenders pathways for decay of the haloferryl intermediate that do not result in l-Lys chlorination, especially at low chloride concentrations; one identified pathway involves oxidation of glycerol. The mechanistic data imply that (i) BesD may have evolved from a hydroxylase ancestor either relatively recently or under weak selective pressure for efficient chlorination and (ii) that acquisition of its activity may have involved the emergence of linkage between l-Lys binding and chloride coordination following loss of the anionic protein-carboxylate iron ligand present in extant hydroxylases.

## Introduction

The selective functionalization of aliphatic carbon centers has been intensively pursued by synthetic chemists but has not yet been routinely realized. Iron-dependent enzymes can install a variety of useful functional groups with high chemo-, regio- and stereo-selectivity.^1–4^ They could potentially replace synthetic transformations that currently require toxic and environmentally harmful reagents with more efficient, greener reactions. Moreover, the degree of control they impart could drastically simplify known synthetic processes by enabling late-stage coupling of advanced intermediates, a capability that could revolutionize production of pharmaceuticals.^5,6^

A large and particularly versatile class of non-heme iron enzymes, the Fe(II)- and 2-oxoglutarate-dependent (Fe/2OG) oxygenases, promote a wide array of outcomes that include hydroxylation,^7^ desaturation,^8^ hetero- and carbo-cyclization,^9^ epimerization,^10^ fragmentation,^11,12^ and halogenation reactions.^13–16^ They perform this range of chemical feats through the substrate-triggered activation of dioxygen at a conserved Fe(II) cofactor, which leads to oxidative decarboxylation of 2OG and formation of an oxoiron(IV) (Fe^IV^=O, ferryl) intermediate.^17–19^ This potently oxidizing high-valent complex cleaves an unactivated C-H bond of the substrate by hydrogen atom (H•) transfer (HAT), generating an intermediate containing a hydroxoiron(III) complex and carbon-centered radical. In the majority of cases, this common intermediate acts as the branch point for the different subclasses of Fe/2OG oxygenases; its fate dictates the reaction outcome. In the most extensively studied subclass, the hydroxylases, the oxygen derived from the ferryl complex is attacked by the radical to form a new C–O bond in a step conventionally referred to as oxygen rebound.^20^ In the Fe/2OG halogenase subclass, a *cis*-coordinated halide, chloride or bromide, is transferred to the carbon-centered radical as a neutral halogen atom (Scheme 1).^21–23^

**Scheme 1:**
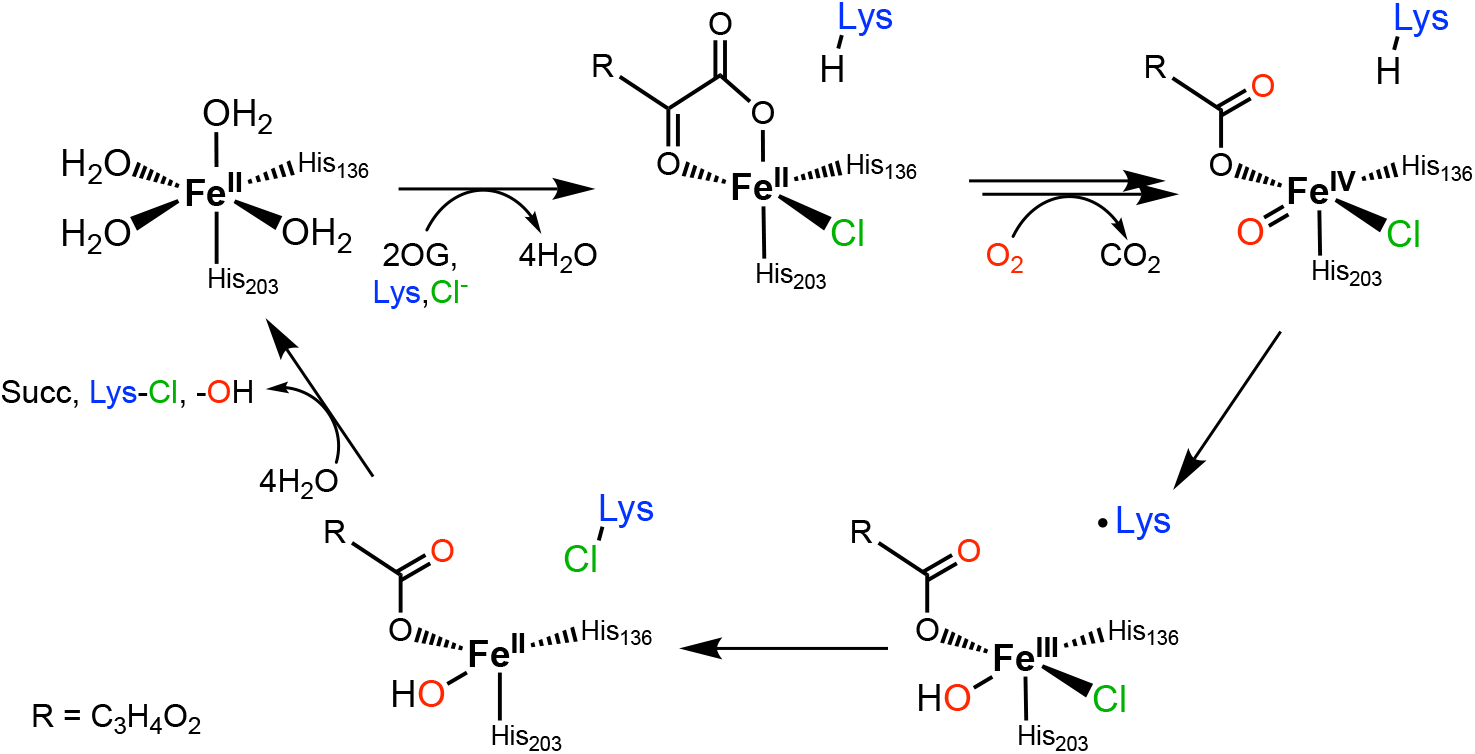
Abbreviated mechanism of haloferryl intermediate formation and ensuing halogenation of l-lysine (at C4) in BesD.

In the halogenases, the disposition of the substrate relative to the *cis*-haloferryl complex has emerged as crucial to ensuring halogen coupling in preference to oxygen rebound, which can be very facile.^17,24^ Studies on the halogenases SyrB2 from the syringomycin E biosynthetic pathway in *Pseudomonas syringae* pv. syringae B301D and WelO5 from the welwitindolinone pathway in *Hapalosiphon welwitschii* UTEX B1830 were instrumental in this recognition.^14,23,25^ However, barriers created by the complexity and/or instability of their substrates have limited deeper insight into how the proper disposition is achieved. SyrB2 natively chlorinates l-threonine linked by a labile thioester to a large carrier protein, SyrB1, while WelO5 chlorinates an indole alkaloid, 12-*epi*-fischerindole U, that is difficult to obtain either naturally or synthetically.^13–15,26^ Recently, a new group of Fe/2OG halogenases with simple amino acid substrates (l-lysine, l-ornithine, l-leucine, l-isoleucine, and l-norleucine) was discovered.^16,27^ One of these halogenases, BesD from *Streptomyces lavenduligriseus*, catalyzes the initial reaction in the biosynthetic pathway to the acetylenic amino acid, <u>β</u>-<u>e</u>thynyl<u>s</u>erine. BesD chlorinates l-lysine at C4 with *R* stereochemistry. BesC fragments the BesD product to l-chloroallylglycine, and BesB then uses the chlorine installed by BesD as the activated leaving group for the final alkyne-forming elimination. BesD provided the first example of an Fe/2OG halogenase that modifies a free amino acid without a carrier protein, thus simplifying kinetic, mechanistic, and structural studies aimed at addressing how oxygen rebound is averted to allow halogen transfer to prevail.

In this study, we set out to assess two issues – the importance of substrate-intermediate disposition in control of the BesD reaction outcome and how the proper disposition is attained – by structurally characterizing vanadyl complexes mimicking the key haloferryl intermediate. We initially found that formation of homogeneous complexes was disfavored by a surprisingly low affinity of the enzyme for vanadyl and strong thermodynamic linkage in binding of the high-valent metal with the halide and l-Lys substrates. We therefore asked whether such thermodynamic linkage extends to functional forms of the enzyme, either its stable Fe(II) states or its reactive Fe(IV)-harboring intermediate. Here, we report dissection of this unusual linkage between the anion and substrate binding in BesD, which has not previously been described in other halogenases. This synergistic binding governs not only the formation of the Fe^II^ quinary complex that activates dioxygen but also the fate of the [*cis*-Cl-Fe^IV^=O] intermediate, as prime substrate and anion can dissociate and re-bind even in this reactive state. This substrate lability engenders pathways for unproductive decay of the reactive species, thus diminishing the efficiency of the enzyme. Through a comprehensive dissection, we identified four different instances of synergistic binding between anion and substrate/product in the BesD halogenase pathway and mapped them onto a kinetic mechanism of catalysis (**Scheme 2**).

**Scheme 2.**
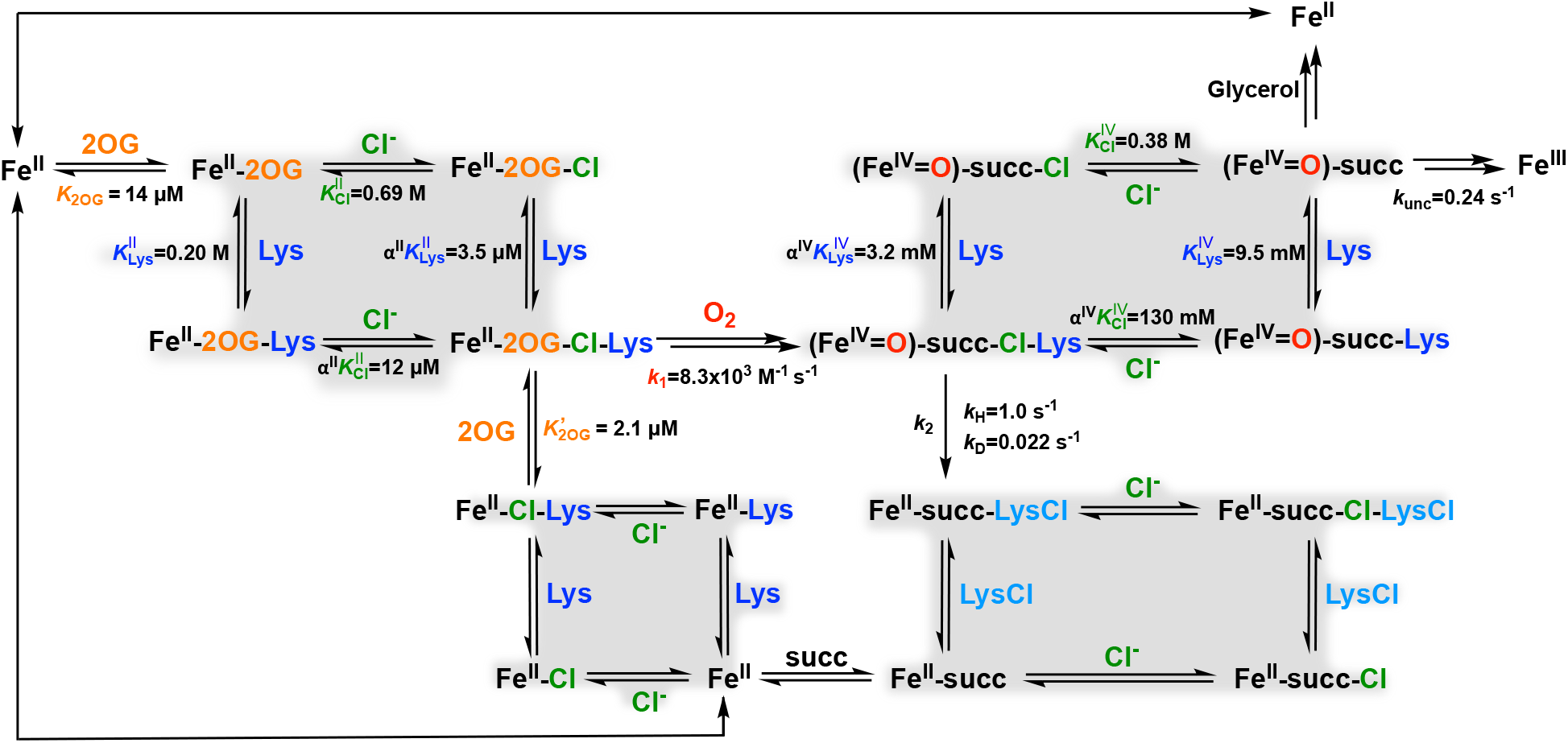
Proposed Kinetic Mechanism for BesD. Cooperative binding cycles are highlighted in gray.

## Results and Discussion

### Cooperative Binding of l-lysine with Anions to BesD

We initially investigated the binding/coordination of anions, chloride, bromide, and azide, to the BesD•Fe^II^•2OG complex by monitoring the resultant shift in the visible absorption band of the complex at ∼ 500 nm arising from the Fe^II^→2OG metal-to-ligand charge-transfer (MLCT) transition that is generally seen in members of this enzyme superfamily. This shift was previously demonstrated in SyrB2, as the addition of Cl^−^ or another anion to the SyrB2•Fe(II)•2OG complex led to hyper- and bathochromic shifts in the MLCT band.^28^ A titration with Cl^−^ revealed a dissociation equilibrium constant, *K*_D_, of 50 µM. In an analogous titration of the BesD•Fe^II^•2OG complex, only modest changes were observed at Cl^−^ concentrations < 200 mM (**Figure 1A**), and saturation of the spectral change required concentrations in the molar range (**Figure 1B & 1C**). Fitting the Δ*A*_520_-versus-[Cl^−^] data (**Figure 1C**) gave a surprisingly high dissociation constant 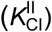 of 0.69 ± 0.07 M. The modest affinity of the BesD•Fe^II^•2OG complex for Cl^−^ was mirrored in titrations with other anions, which gave *K*_D_ values of 2.0 ± 0.3 M for Br^−^ and 0.19 ± 0.01 M for N3– (**Figure S1**). On the basis of this measured 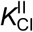, BesD would be only fractionally saturated by its anion co-substrate under the physiological growth conditions of the native organism (∼120 mM NaCl).^29^

**Figure 1.**
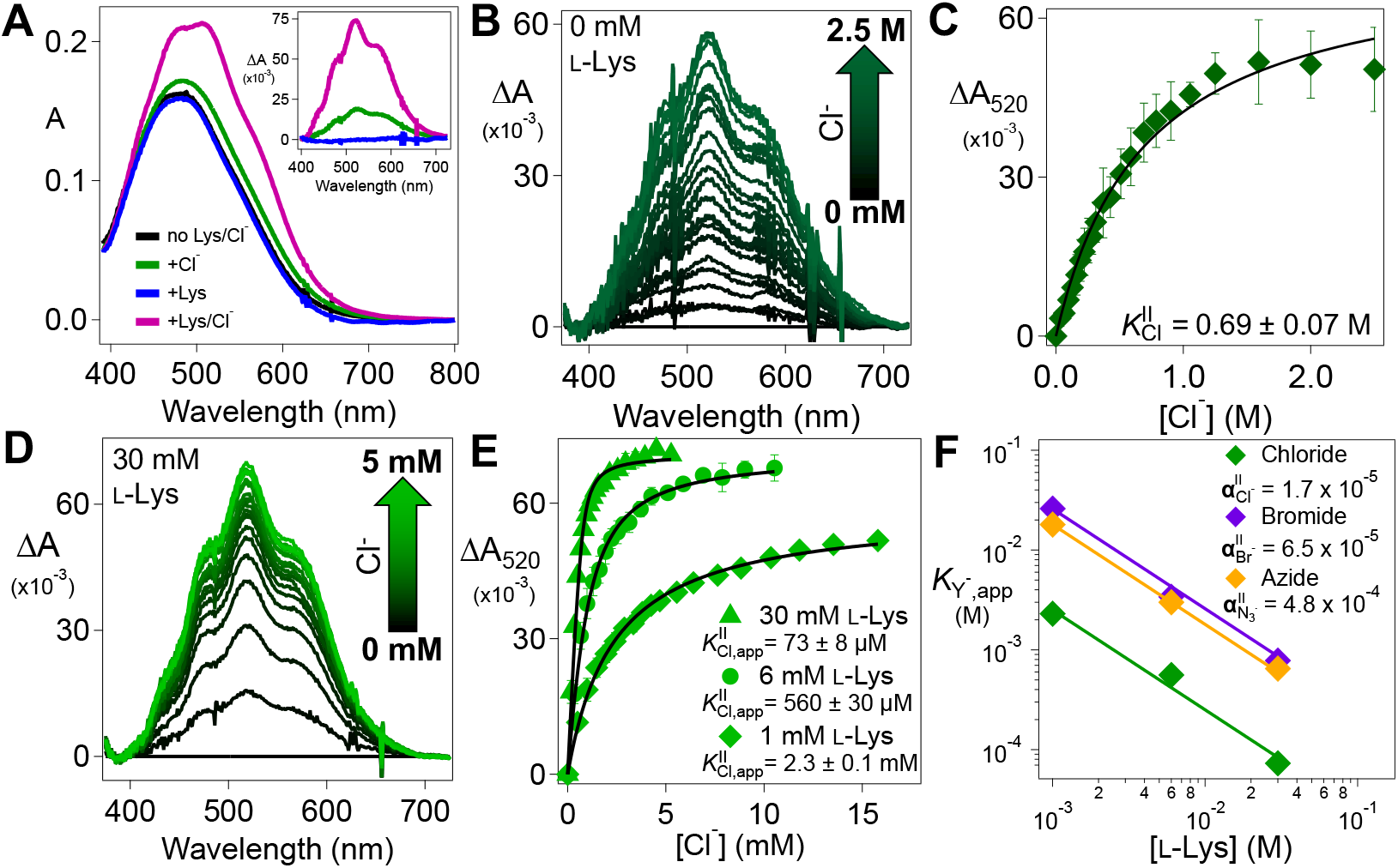
(***A***) UV-visible absorption spectra of a solution of 1.0 mM BesD, 0.72 mM Fe^II^, and 5 mM 2OG in the absence of l-Lys and NaCl (*black*), the presence of just 200 mM NaCl (*green*), the presence of just 5 mM l-Lys (*blue*), and the presence of both 5 mM l-Lys and 200 mM NaCl (*pink*). (*inset*) Difference spectra obtained by subtracting the black spectrum from the others. (***B***) Difference spectra obtained after addition of aliquots of a concentrated NaCl solution to a solution of the BesD•Fe^II^•2OG complex in the absence of l-Lys. (***C***) *ΔA*_520_-versus-[Cl^−^] titration curve (n=3). 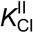 was determined by fitting the data by the hyperbolic equation for binding. (***D***) Difference spectra from a similar titration but in the presence of 30 mM l-Lys. (***E***) *ΔA*_520_-versus-[Cl^−^] titration curves from experiments carried out in the presence of 1 mM (*diamonds*), 6 mM (*circles*), and 30 mM l-Lys (*triangles*) (n=3 for each). 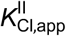 was determined by fitting the data to either the hyperbolic or the quadratic curve for binding, as appropriate. (***F***) Cooperativity plot for K_Y_-_,app_ as a function of total [l-Lys] for Cl^−^ (*green*), Br^−^ (*purple*), and N_3_ ^−^ (*orange*). The data were fit by equation 1 with constants 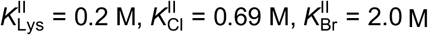, and equation 1 with constants 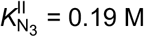.

However, inclusion of a sufficient concentration of the prime substrate, l-lysine, in the BesD•Fe(II)•2OG solution was found to diminish the [Cl^−^] needed to elicit the MLCT shift (**Figure 1A**). Chloride titrations with l-lysine in the 1-30 mM range yielded values of 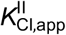 having an inverse relationship to the [l-Lys]. In other words, increasing [l-Lys] increases the apparent affinity of BesD for Cl^−^. The apparent affinities for Br^−^ and N3– are similarly affected by [l-Lys] (**Figure S2**).

To demonstrate the reciprocal effect of Cl^−^ on l-Lys affinity, we used microscale thermophoresis (MST) to monitor l-Lys association to the BesD•Fe^II^•2OG complex. l-Lys has a relatively high dissociation equilibrium constant, 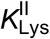, of 0.20 ± 0.06 M in the absence of Cl^−^ (**Figure S3**). Similarly to the effect of [l-Lys] on the Cl^−^ titrations, the presence of 6 mM Cl^−^ was found to decrease the apparent *K*_D_, 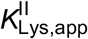, of l-Lys to 0.40 ± 0.14 mM (**Figure S3**). Along with the Cl^−^ titrations, this increase in apparent affinity for l-Lys with increasing [Cl^−^] implies a strong heterotropic cooperativity between l-Lys and Cl^−^ binding by BesD. The large magnitude of the cooperativity term (see below) presents a challenge to deduction of their binding order. As such, the assembly of the complex is best modeled by a square equilibrium scheme (Scheme 2, *upper left*). From this model, we derived expressions (see *SI* text) for the cooperativity coefficient in the Fe^II^-containing complex, **ΔII**, in terms of the apparent dissociation constants, 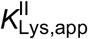 and 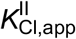, obtained from the individual l-Lys and Cl^−^ titrations in the presence of varying concentrations of the other substrate (**Figures 1E** and **S3**):

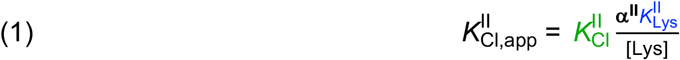

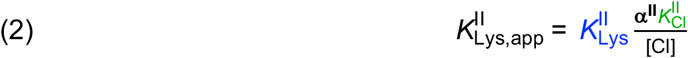

α**Δ^II^** is the factor by which the binding of one substrate changes the effective dissociation constant for the other. It can be found (independently) from either expression and the data from the optically- and MST-monitored titrations described above. The Cl^−^ titrations yielded an estimate of α**^II^** = 1.7×10^−5^ ± 0.6×10^−5^, and the l-Lys MST titrations gave **Δ^II^** = 1.7×10^−5^ ± 0.8×10^−5^. The excellent agreement establishes a strong cooperativity in binding of the anion and prime substrate in BesD. In essence, binding of l-Lys increases affinity for Cl^−^ by almost 10^5^ and vice versa. Using the same methodology, we found the non-native anions Br^-^ and N_3_-to be only slightly less cooperative with the prime substrate (α**Δ^II^** = 6.5×10^−5^ and 4.8×10^−4^, respectively; **Figure 1F**). The preferential affinity for the native anion, Cl^−^, is congruent with the prior anion-binding studies on SyrB2.^28^ Similarly, we evaluated two l-Lys analogs, D-Lys and l-ornithine, and found their effective dissociation constants with chloride bound (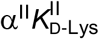 and 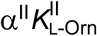) to be ∼ 3- and 20-fold greater, respectively, than that of l-Lys (**Figure S4**), comporting with the fact that l-Lys is the native prime substrate of BesD.

### Evidence for a Preferred Order of Substrate Binding in Assembly of the O_2_-reactive Quinary Complex

BesD must bind its Fe(II) cofactor and three co-substrates (2OG, prime substrate, and anion) before dioxygen can add. Assembly of the O_2_-reactive BesD•Fe^II^•2OG•l-Lys•Cl complex (hereafter the quinary complex) could have a preferred or even obligatory binding order. By monitoring the ∼ 500-nm MLCT band, we found that binding of 2OG to the BesD•Fe^II^ binary complex is associated with a *K*_D_, *K*2OG, of 14 ± 5 µM. In the presence of l-Lys and Cl^−^ under saturation conditions (40 mM l-Lys, 1 M Cl^−^), the *K*_D_ for 2OG, denoted 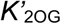, decreases to 2.1 ± 1.0 µM, implying only a modestly increased affinity for 2OG in the fully assembled quinary complex (**Figure S5**). This small difference contrasts with the strong linkage between l-Lys and Cl^−^ and implies that 2OG binding is nearly independent.

Previous investigations into the order of substrate association in the Fe/2OG hydroxylase, TauD, showed that addition of 2OG to the TauD•Fe^II^ complex is fast but can be slowed by prior incubation of the prime substrate, taurine, with the enzyme.^19^ This effect of taurine was not observed when the TauD•Fe^II^ complex was mixed simultaneously with both taurine and 2OG, indicating a preference for 2OG binding before the prime substrate. We performed an analogous order-of-mixing kinetic analysis of how BesD acquires its *three* substrates.

To determine if 2OG binding to the Fe^II^ center can be slowed by complexation with l-Lys and/or Cl^−^, we tested all possible permutations of mixing order (**Figure S6**). Inclusion of either chloride or l-Lys in the BesD•Fe^II^ solution that was mixed with 2OG resulted in only minor amplitude changes in the *βA*520 signal, with no obvious change in the formation kinetics (**Figure 2**, *blue, purple, and green traces*). By contrast, inclusion of both l-Lys and Cl^-^ with BesD•Fe^II^ slowed 2OG association by more than a factor of 100 (*red trace*). Similarly to TauD, the presence of prime substrate in the active site interferes with 2OG association, but, in BesD, the linkage between l-Lys and Cl^−^ binding requires the presence of both to form the complex and realize the interference. This result also implies that l-Lys and Cl^−^ can bind to the BesD•Fe^II^ complex in the absence of 2OG, with the synergy (heterotropic cooperativity) shown more directly for their binding to BesD•Fe^II^•2OG. The slower binding of 2OG is likely a result of the C-terminal lid-loop region, common to Fe/2OG enzymes, closing off the active site as a result of prime substrate docking. This phenomenon was recently observed by comparison of structures of a related l-lysine 5-chlorinase, HalB, either lacking or containing l-Lys.^30^

**Figure 2.**
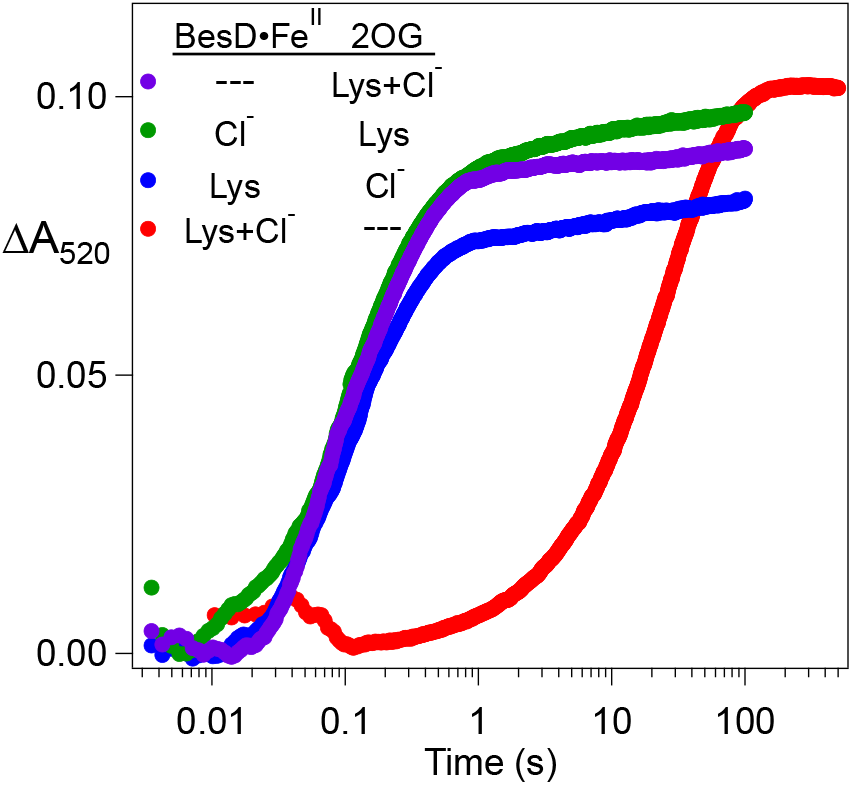
Kinetics of 2OG binding to BesD•Fe^II^ (and higher complexes). Equal volumes of anoxic solutions containing (1) 1.1 mM BesD and 0.8 mM Fe^II^ and (2) 2.4 mM 2OG were rapidly mixed at 5 °C, and development of the MLCT band at 520 nm was monitored. The color-coded table indicates the other substrates added to solutions 1 and 2 in the individual experiments. When present, l-Lys was at 80 mM and Cl^−^ was at 2 M (before mixing).

### Substrate-triggered Activation of O_2_ by BesD•Fe^II^•2OG•Cl^−^ Generates the Chloroferryl Intermediate

Ferryl intermediates have been directly demonstrated in Fe/2OG enzymes belonging to multiple subclasses. In each case, O_2_ adds last to the enzyme complex with Fe^II^ and all other substrates bound. Partial complexes lacking the prime substrate or, for the halogenases SyrB2 and HctB, a coordinated anion (natively, halide) have been shown to react much less rapidly with O_2_, and, in these cases, accumulation of the ferryl complex has not been observed.^28,31^ We have termed this activation of the cofactor for reaction with O_2_ “substrate triggering.” BesD is similarly substrate-triggered for reaction with O_2_ to form a transient UV-absorbing ferryl complex. Mixing of the pre-formed BesD•Fe^II^•2OG•l-Lys•Cl^−^ quinary complex with a solution containing O_2_ resulted in development and decay of the characteristic absorption feature at ∼ 320 nm (**Figure 3A**, *red line*). By contrast, if we omitted either Cl^−^ or l-Lys from the protein reactant solution, we saw only slow development of UV absorption characteristic of oxidation of the Fe^II^ cofactor to Fe^III^ (*black, green, and blue lines*).

**Figure 3.**
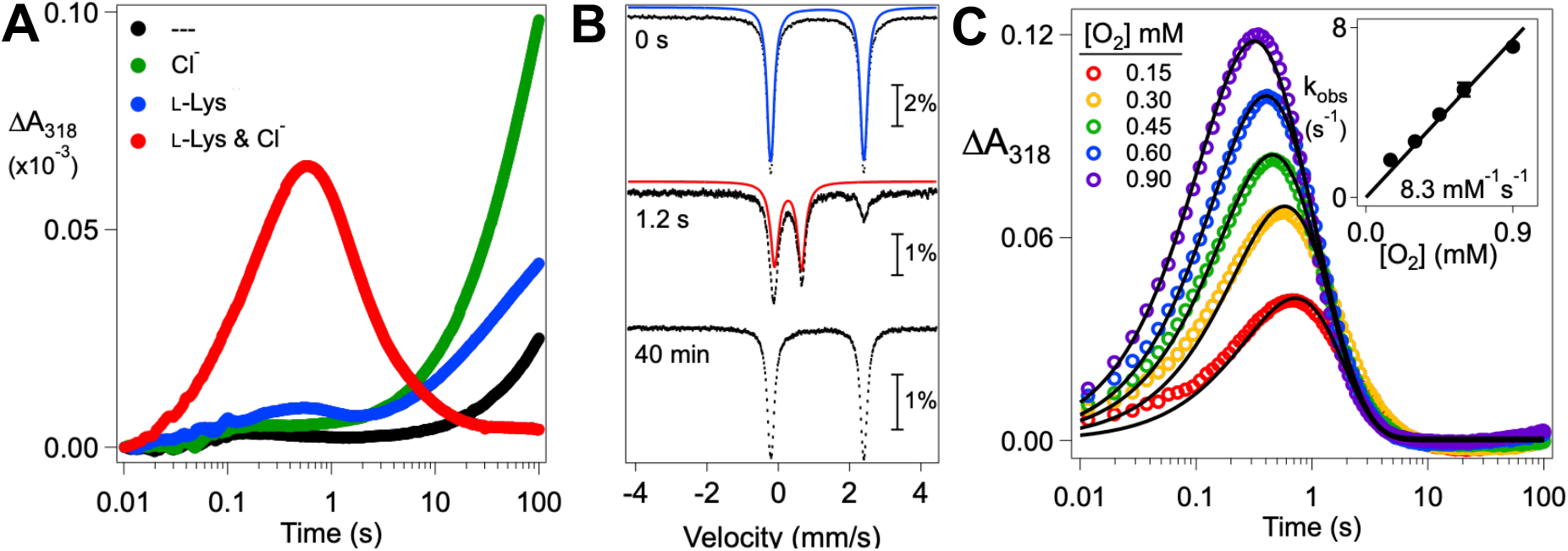
Substrate-triggered, O_2_-dependent accumulation of the chloroferryl intermediate in BesD. (***A***) Change in absorbance at 318 nm (*ΔA*_318_) as a function of time after an anoxic solution containing 0.55 mM BesD, 0.4 mM Fe^II^, and 5 mM 2OG in the absence or presence of 3 mM l-Lys and/or 200 mM Cl^−^ was mixed at 5 °C with an equal volume of air-saturated buffer (giving ∼ 0.18 mM [O_2_] after mixing). (***B***) Mössbauer spectra (4.2 K/0 mT) of samples prepared either by freezing an anoxic solution containing 1.8 mM BesD, 1.5 mM ^57^Fe^II^, 16 mM 2OG, 10 mM *d*_4_-l-Lys and 100 mM NaCl (*upper spectrum*) or by mixing this solution at 5°C with an equal volume of O_2_-saturated buffer (giving ∼0.9 mM [O_2_] after mixing) and freezing at the reaction times indicated in the figure (*middle and lower spectra*). Solid lines are simulations of the spectra of Fe^II^ (*blue*) and Fe^IV^ (*red*) components. The isomer shift and quadrupole splitting parameters of the Fe^II^ and Fe^IV^ species are δ_Fe(II)_ = 1.09 mm/s, ΔE_Q,Fe(II)_ = 2.62 mm/s and δ_Fe(IV)_ = 0.27 mm/s, ΔE_Q,Fe(IV)_ = 0.73 mm/s. (***C***) *ΔA*_318_-versus-time traces after an anoxic solution containing 0.28 mM BesD, 0.2 mM Fe^II^, 0.2 mM 2OG (ensuring a single turnover), 800 mM NaCl, and 10 mM l-Lys was rapidly mixed at 5 °C with an equal volume of O_2_-containing buffer delivering the final concentrations shown in the figure. Traces were fit by the equation describing absorbance from an intermediate that forms and decays in irreversible first-order reactions from and to transparent reactant and product states, as described in the *SI* text. In the global analysis, the effective first-order formation rate constants (*k*_obs_) were allowed to vary with [O_2_], but the first-order decay rate constant was required to be identical for each fit. (*Inset*) The means of the best-fit *k*_obs_ values versus final [O_2_], with linear fit through the origin. The error bars reflect the range in two independent trials.

We confirmed the identity of the transient UV-absorbing complex by freeze-quench Mössbauer spectroscopy. We used deuterated l-lysine, 4,4,5,5,-[^2^H4]-l-Lys (*d*4-l-Lys), to leverage the expected large substrate deuterium kinetic isotope effect (D-KIE) on HAT (which is common to ferryl intermediates in this class of enzymes) for greater accumulation of the intermediate.^15,17,18^ The quadrupole doublet associated with the reactant complex (**Figure 2B**, *top, blue line*) has parameters (δ = 1.09 mm/s, ΔEQ = 2.62) typical of high-spin Fe^II^.^32^ In the spectrum of a freeze-quenched sample prepared by reacting this complex with O_2_ for 1.2 s (the time of maximum accumulation of the intermediate; see below), the major component is a quadrupole doublet (**Figure 2B**, *middle, red line*) with parameters (δ = 0.27 mm/s, ΔEQ = 0.73 mm/s) typical of high-spin ferryl complexes detected in other Fe/2OG enzymes.^33^ Consistent with the assignment as a ferryl intermediate, its features are not present in the spectrum of a sample allowed to react for 40 min (**Figure 2B**, *bottom*). The Mössbauer parameters of this single ferryl species in BesD are similar to those of one of the two chloroferryl species observed in the halogenase SyrB2 (δ = 0.23 mm/s and ΔEQ = 0.76 mm/s).^15^

Although the ferryl complex of an Fe/2OG oxygenase is not the first intermediate in the reaction of its enzyme•substrates complex with O_2_, the complex often forms with kinetics characteristic of a biomolecular reaction, implying that no ferryl precursor accumulates.^19^ To test for this possibility in BesD, the dependence of development of the UV-absorption feature of the chloroferryl complex on the concentration of O_2_ was delineated under single-turnover conditions (**Figure 3C**). The *ΔA*318-versus-time traces – specifically their formation phases – show the anticipated dependence on [O2]. Best-fit values for the effective first-order formation rate constant, *k*obs, show a linear dependence on [O2] and give a second-order rate constant of 8.3 ± 0.3 mM^-1^ s^-1^ (**Figure 3C**, *inset*). The linked decay rate constant determined by the global fitting of two trials of five oxygen concentrations is 1.053 ± 0.003 s^-1^.

### Unproductive Decay of the Chloroferryl Intermediate Diminishes the Observed Substrate D-KIE

For several Fe/2OG enzymes, the role of the ferryl complex in abstracting H• from the substrate has been established by demonstration of a large, normal substrate D-KIE on decay of the intermediate. Intrinsic D-KIEs on the HAT step of 30-60 have been determined.^8,15,17,18,34^ However, the large intrinsic D-KIE can result in engagement of alternative pathways for ferryl intermediate decay, including abstraction of H• from a different, protium-bearing carbon of the substrate, unproductive decay by one-electron reduction, or both. By affording an escape route not requiring deuterium transfer, this situation can mask a large intrinsic D-KIE on the HAT step and make the observed effect on ferryl intermediate decay much smaller.^34,35^ Initial characterization of the Bes biosynthetic pathway established that C4 of l-Lys is the site of halogenation by BesD. We therefore used commercially available 4,4,5,5-[^2^H4]-l-Lys (*d*4-l-Lys) to test for the expected D-KIE on ferryl intermediate decay.^16,27^ Initial experiments with substrate concentrations in the range used in prior studies of other halogenase (1.5 mM prime substrate and 40 mM NaCl) gave an unexpectedly small *observed* D-KIE of 5-6 (**Figure 4A**, *dashed traces*). To rule out the possibility that the presence of deuterium at C4 (and C5) redirects the ferryl complex to abstract H• from a different site (C2, 3, or 6), we also tested 2,3,3,4,4,5,5,6,6-[^2^H9]-l-Lys (*d*9-l-Lys) as substrate. Use of this substrate did not detectably change the reaction kinetics relative to those seen with the *d*4-l-Lys (**Figure S7**), implying that the BesD ferryl complex targets C4 (or C4 and C5) with high specificity. Precedent suggests that the residual absorbance at long times in the *βA*318-versus-time traces from the reactions with these deuterated substrates reflects net conversion of the cofactor from the nearly transparent Fe^II^ state to the UV-absorbing Fe^III^ state via the aforementioned one-electron reduction of the ferryl complex.^15,22^ Analysis by liquid chromatography-mass spectrometry (LC-MS) of the products from BesD reactions carried out under similar (single-turnover) conditions with 20 mM chloride revealed that only ∼ 9% as much chlorinated product forms from the *d*4-l-Lys isotopolog as forms from unlabeled l-Lys. (**Figure 4B**, and **4C**, *blue and red bars*). The amount of succinate produced in ferryl formation as a byproduct of 2OG decarboxylation (*light blue and peach bars*) indicates that, as expected, substrate deuteration does not impact the quantity of ferryl intermediate formed (at any [Cl^-^]). These observations are consistent with a process involving unproductive decay of the chloroferryl intermediate.

**Figure 4.**
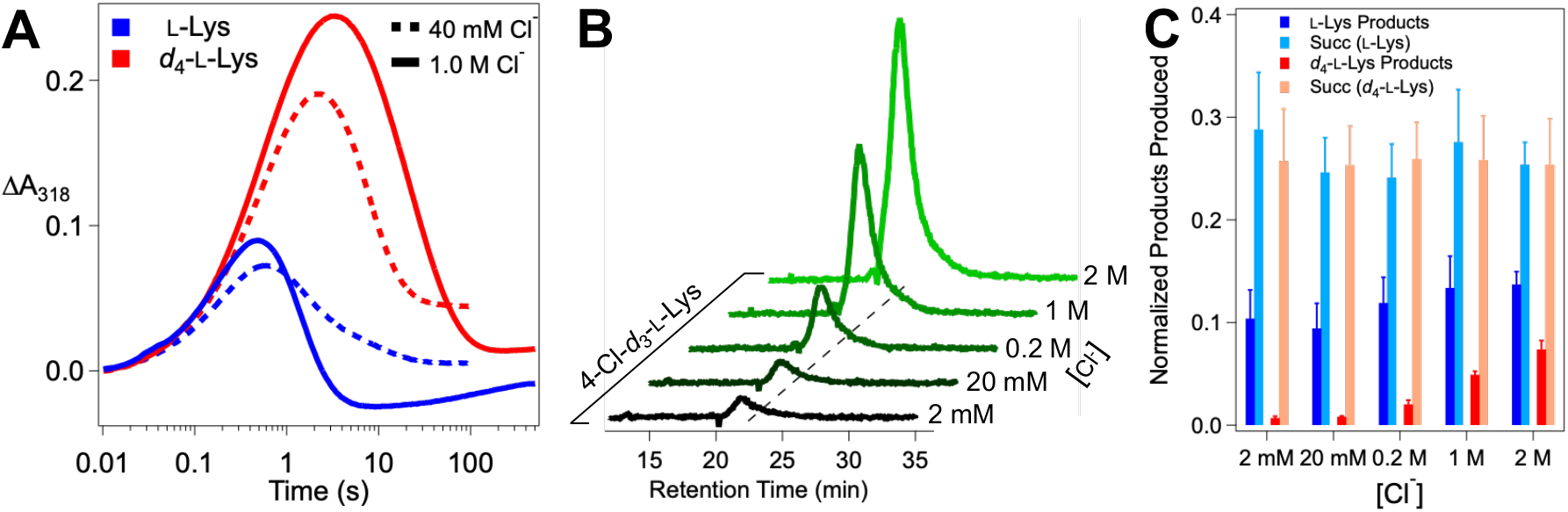
(***A***) Kinetics of formation and decay of the BesD chloroferryl intermediate in reactions with l-Lys (*blue*) and *d*_4_-l-Lys (*red*) at high (*solid lines*) and low (*dashed lines*) [Cl^−^] as monitored by its absorbance at 318 nm. An anoxic solution containing 0.55 mM BesD, 0.4 mM Fe^II^, 5 mM 2OG, 10 mM l-Lys/*d*_4_-l-Lys, and either 80 mM or 2.0 M NaCl was mixed at 5 °C with an equal volume of air-saturated buffer (∼0.36 mM [O_2_]). (***B***) Single-ion chromatograms monitoring 4-^35^Cl-*d*_3_-l-Lys (184 m/z) produced from single-turnover assays at varying [Cl^−^]. (***C***) Normalized products by single-turnover assays with l-Lys and *d*_4_-l-Lys at varying concentrations of NaCl. l-Lys/*d*_4_-l-Lys derived product peak areas are a combination of the detected 4-Cl-l-Lys (^35^Cl-& ^37^Cl-) and its decomposition products: 4-OH-l-Lys and 2-amino-4-propylamine-γ-lactone/2-amino-4-hydroxy-ε-lactam (see Figure S8 for decomposition pathway). The product peak areas were normalized to an internal standard of the opposite isotopolog added after the reactions were quenched. Succinate produced was normalized to an internal standard of *d*_4_-succinate added after quenching. For reaction conditions, see the *SI Text*.

Use of higher chloride concentrations (e.g., 1.0 M) was found to increase the amplitude of the *βA*318 transient and impact the decay phase (**Figures 4A**, *solid traces*). For the reaction with the protium-bearing l-Lys, the decay phase becomes just slightly *faster* at the higher [Cl^−^] (*blue traces*), whereas for the reaction with *d*4-l-Lys, it becomes markedly *slower* (*red traces*). These opposing effects cause the observed D-KIE on ferryl decay to increase from ∼ 7 at 40 mM Cl^−^ (0.89 s^-1^ / 0.13 s^-1^) to ∼ 32 at 1.0 M Cl^−^ (1.2 s^-1^ / 0.038 s^-1^). The presence of 1.0 M [Cl^−^] also allows UV absorbance to decay almost completely (i.e., for *βA*318 to approach zero at long reaction times), suggesting that the high [Cl^−^] suppresses unproductive ferryl decay (**Figure 4A**). Importantly, the yield of the 4-Cl-4,5,5-*d*3-l-Lys product from the *d*4-l-Lys substrate also increases with increasing [Cl^−^], whereas succinate production is not affected (**Figure 4B, C**). These observations imply that another reaction competes with slow abstraction of deuterium from *d*4-l-Lys and that the partition ratio is affected by [Cl^−^], with higher concentrations favoring the productive branch. Given the unexpected lability of the anion and prime substrate in the BesD reactant state and the ability of increasing [Cl^−^] to suppress the unproductive pathway, we posited that it might be initiated by chloride and possibly l-lysine dissociation *from the chloroferryl intermediate state*. If so, its suppression by high [Cl^−^] would reflect rebinding of the substrate(s) in competition with the next step in the unproductive pathway, such as reduction of the Fe^IV^. Although we viewed the idea of reversible binding of substrates in an intermediate state of sufficient potency to cleave an unactivated C–H bond as surprising, we did recently demonstrate such lability in the substrate-triggered peroxodiiron(III) intermediate state in the diiron enzyme BesC, which fragments 4-Cl-l-Lys and l-Lys – putatively by abstracting hydrogen from C4 – in the β-ethynylserine biosynthetic pathway.^36^

### Synergistic Binding of Cl^−^ and l-Lys in the Chloroferryl Intermediate State

In light of the demonstrated binding synergy between the anion and prime substrate in the Fe^II^ state of BesD, we further evaluated the possibility of reversible substrate association in the intermediate state by testing for an interplay between the Cl^−^ and l-Lys concentrations in the kinetics of decay of the chloroferryl intermediate and its partitioning between productive and unproductive pathways. Indeed, we found that an increasing concentration of either substrate at a fixed concentration of the other increases the lifetime of the chloroferryl intermediate (**Figure 5A, B**), thus confirming an analogous synergistic interaction between prime substrate and anion in the chloroferryl intermediate state. The expectation that ferryl complexes lacking a substrate can decay only unproductively connects the kinetic effects of their concentrations to their reversible binding (**Figure 5C**). This model yields an analytical expression (**Equation 3**) for the rate constant for decay of the ferryl intermediate (*K*_D_ecay) in terms of the substrate concentrations, their equilibrium dissociation constants (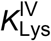 and 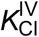), their cooperativity coefficient in the high-valent state (**Δ**α**IV**), the rate constant for productive decay via deuterium abstraction (*K*_D_), and the rate constant for unproductive decay following substrate dissociation (*k*unc):

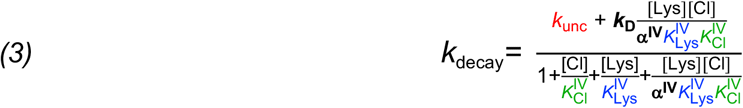

A derivation of this expression is provided in the *SI Text*. We more comprehensively mapped the variation of *K*_D_ecay with [Cl^−^] and [*d*4-l-Lys] via an 11 × 4 ([Cl^−^] x [*d*4-l-Lys]) matrix. We fit the individual *ΔA*318-versus-time traces by Equation 4 to extract values of *K*_D_ecay [the sum of productive (*k*2) and unproductive (*k*3) rate constants] at each combination of [Cl^−^] and [*d*4-l-Lys].

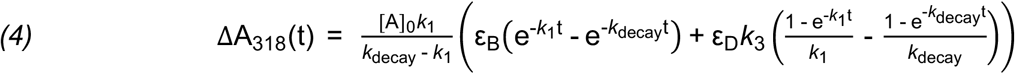

In Eq. 4, [A]0 is the initial limiting reactant concentration (O2 under these conditions), *k*1 is the effective first-order formation rate constant, and εB & εD are the molar absorption coefficients of the ferryl and ferric states, respectively. We then plotted the extracted *K*_D_ecay values as a function of [Cl^−^], giving a family of four curves representing the four different values of [*d*4-l-Lys] (**Figure 5D**). We fit all four replots globally according to Equation 3 by linking the five coefficients, 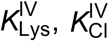, 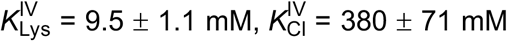, α**ΔIV**, *K*_D_, and *k*unc, and keeping [*d*4-l-Lys] fixed at the known value for each curve. From this analysis, the coefficients converged to values of 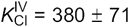, α**ΔIV** = 0.34 ± 0.18, *K*_D_ = 0.022 ± 0.004 s^-1^, and *k*unc = 0.24 ± 0.01 s^-1^. An analysis of the uncertainties in the parameters is provided as **Figure S9**. The rate constant for deuterium transfer to the ferryl complex from this analysis (0.022 s^-1^) can be compared to that for protium transfer (1.035 s^-1^; **Figure 3C**) to yield an intrinsic D-KIE of ∼ 45 on the HAT step, in line with values determined in the reactions of other Fe/2OG enzymes.^8,15,17,18,34^

**Figure 5.**
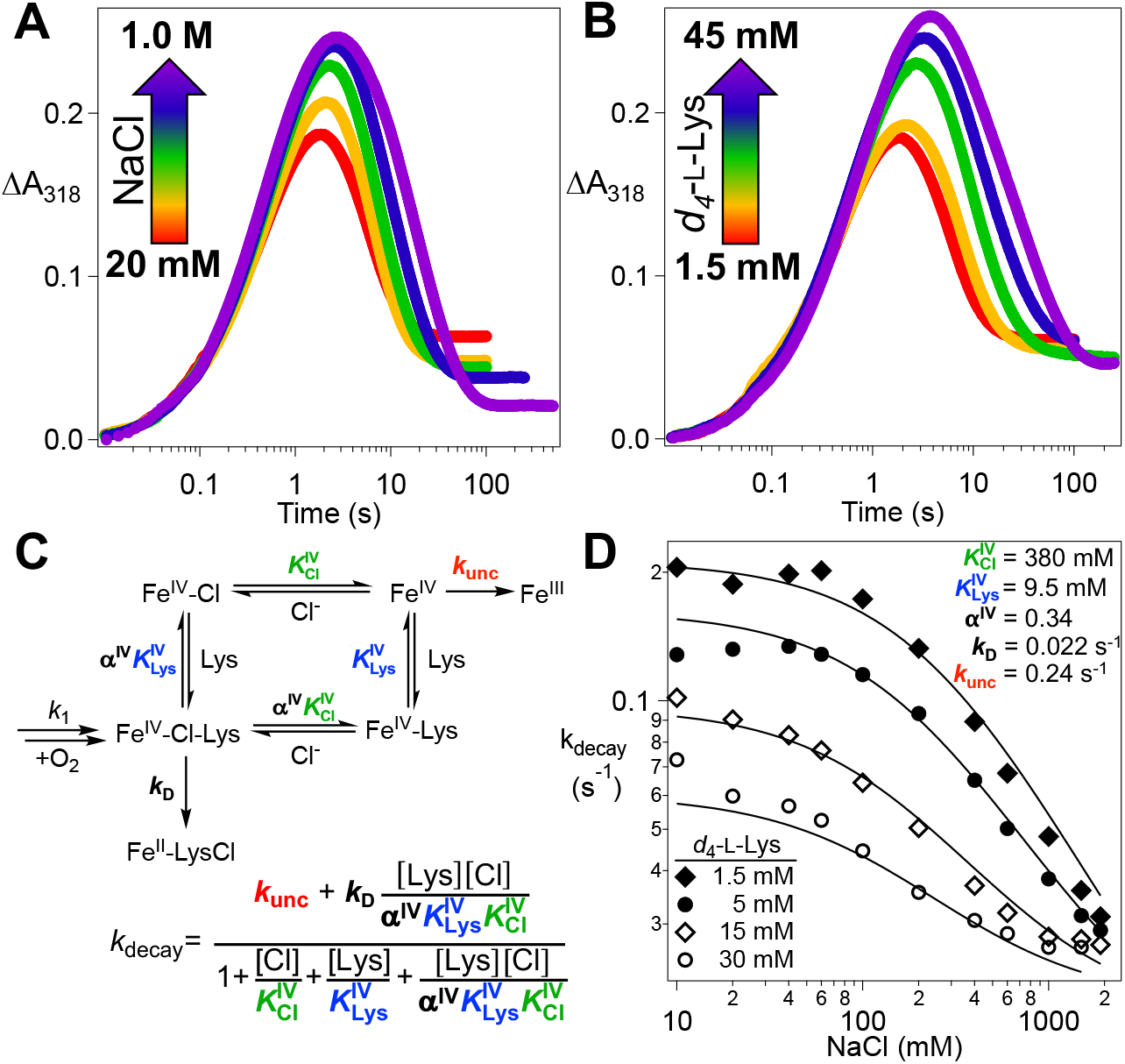
Dependence of the kinetics of the BesD (chloro)ferryl intermediate on the *d*_4_-l-Lys and chloride concentrations and analysis of reversible substrate binding in the high-valent state. (***A***) *ΔA*_318_-versus-time traces after an anoxic solution containing 0.55 mM BesD, 0.4 mM Fe^II^, 5 mM 2OG, 3 mM *d*_4_-l-Lys, and varying NaCl concentration was rapidly mixed at 5 °C with an equal volume of air-saturated buffer (∼ 0.18 mM final [O_2_]). (***B***) Traces from experiments carried out as in ***A*** but with varying *d*_4_-l-Lys and 40 mM NaCl in the protein syringe. The substrate concentrations indicated by the color coding are those after mixing. (***C***) Mechanistic model for the reversible, synergistic binding of l-Lys and chloride in the ferryl state and for its decay via productive and unproductive pathways with the substrates bound and unbound, respectively. (***D***) Observed values of the rate constant for decay of the ferryl state as a function of [Cl^−^] at four values of [*d*_4_-l-Lys]. Traces were globally fit by equation 3.

The dissociation of substrates from the intermediate state is obvious in these data because deuterium transfer is so slow that unproductive decay initiated by dissociation becomes competitive at low substrate concentrations. The underlying phenomenon is also present in reactions with the native protium substrate, but the much faster protium transfer still out-competes dissociation, as demonstrated by the relative insensitivity of product yield to [Cl^−^] (**Figure 4C**). In essence, the lability of the complete chlorinating complex in BesD is countered by a relatively fast HAT step, which may explain why BesD evolved to be more efficient in this step than SyrB2.^15^

The linked substrate binding thermodynamics in BesD change in interesting ways from the resting Fe^II^ state to the reactive Fe^IV^ state. 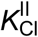 and 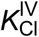 are similar (690 mM vs. 380 mM, respectively), but the affinity for l-Lys increases from 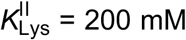 to 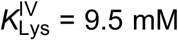. These ∼ 2- and ∼ 20-fold increases in affinity are insufficient to fix the substrates in place, owing to an even more substantial loss of cooperativity, from **Δ^II^** = 1.7 × 10^−5^ to **Δ** α**IV** = 0.34. This 10^4^ loss in cooperativity diminishes complex stability so much that, when the slower deuterium transfer affords time for equilibrium to be approached, unproductive decay pathways become competitive unless substrate concentrations are much higher than are required merely to trigger chloroferryl intermediate formation by driving formation of the reactant quinary complex. This relatively facile uncoupling pathway probably would not have been selected against during the evolution of BesD, for the simple reason that it is engaged to a significant extent only when it competes with a HAT step artificially delayed by the presence of deuterium in the substrate.

### Spectroscopic Evidence for One-Electron Reduction of the Ferryl Complex to Fe^III^ Species

As noted, residual UV absorption at long reaction times seen in the stopped-flow experiments at low substrate concentrations suggests conversion of the reactive Fe^IV^ state to stable Fe^III^ products in competition with on-pathway return of the cofactor to its resting Fe^II^ state. We used Mössbauer spectroscopy to verify this inference. In freeze-quenched samples prepared with either low concentrations (5 mM *d*4-l-Lys, 50 mM Cl^-^) or high concentrations (30 mM *d*4-l-Lys, 1.5 M Cl^-^) of the substrates, conversion of the ferrous reactant complex to the ferryl state is obvious in the 4.2-K/0-mT Mössbauer spectra (**Figure 6A, B**, *compare top and middle spectra*), as shown previously in **Figure 3B**. At the lower substrate concentrations, the intermediate fully decays by 30 s (**Figure 6A**, *bottom*). At the higher concentrations, a significant quantity of the intermediate remains at this reaction time (**Figure 6B**, *bottom*), and complete decay requires ∼ 100 s (**Figure S10**). In addition, more of the intermediate accumulates under the high-[substrates] conditions (69% compared to 46%). These observations closely mirror the trends seen in the stopped-flow absorption experiments, in which higher concentrations of the substrates were seen to increase the amplitude of the transient UV absorption and slow its decay.

**Figure 6.**
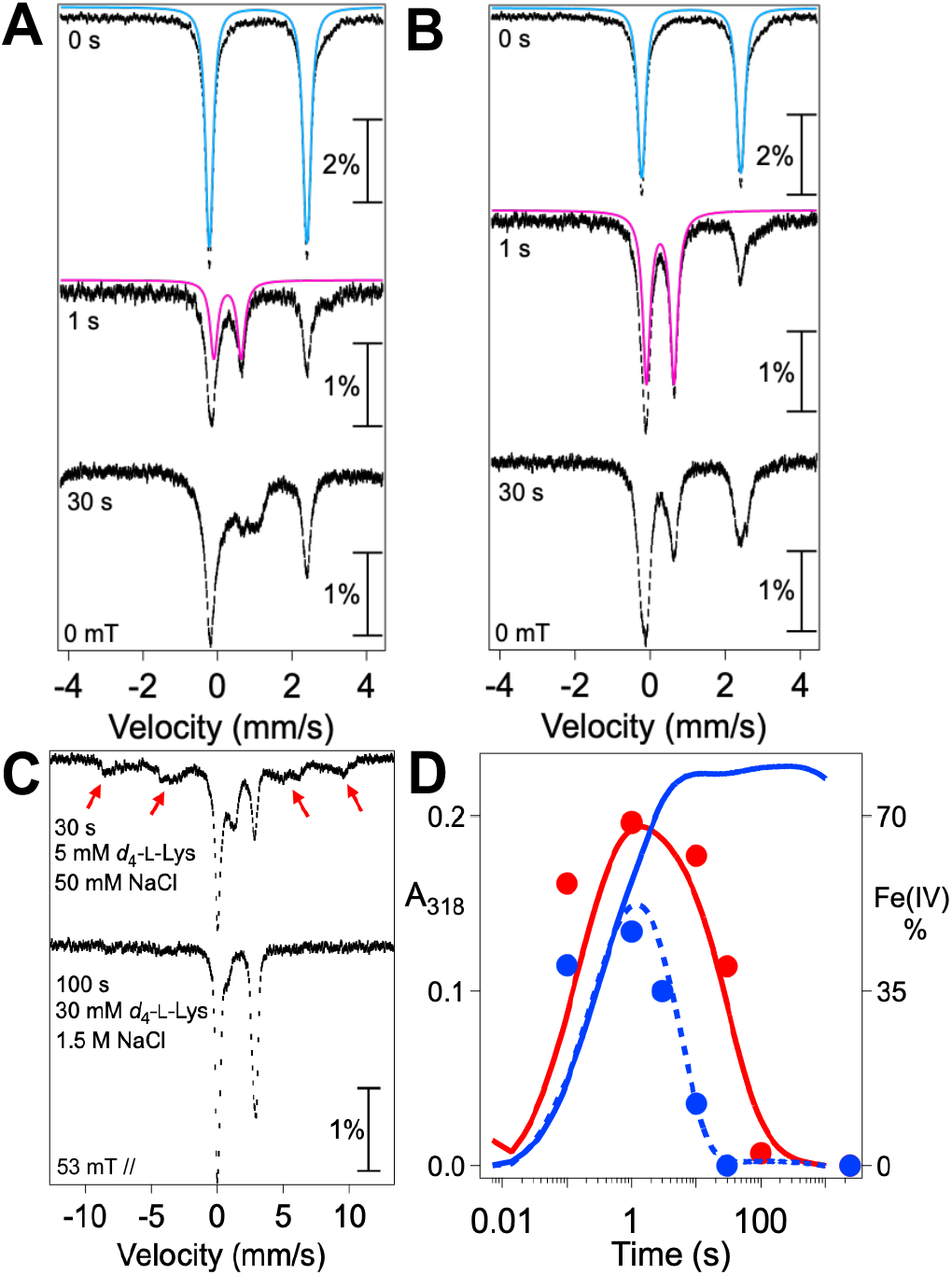
Mössbauer characterization of BesD reactant complexes at high and low [Cl^−^] and [l-Lys] and of their reactions with O_2_. (***A*** and ***B***) Spectra at 4.2 K of samples prepared either by manual freezing of an anoxic solution containing 1.8 mM BesD, 1.5 mM ^57^Fe(II), 16 mM 2OG, and (***A***) 10 mM *d*_4_-l-Lys and 100 mM NaCl or (***B***) 60 mM *d*_4_-l-Lys and 3.0 M NaCl (*top spectra*), or by mixing one of these solutions with an equal volume of O_2_-saturated buffer (∼ 0.9 mM O_2_ after mixing) and freezing at the reaction time indicated in the figure (*middle and bottom spectra*). The reference spectra of the Fe^II^ quinary complex (*blue line*) and ferryl species (*magenta line*) are described in **Figure 3**. (***C***) 4.2 K/53 mT (field parallel to γ-beam) spectra of samples from ***A*** (*top spectrum*) and ***B*** (*bottom spectrum)* after complete decay of the ferryl species. Broad features from the paramagnetic species that develop(s) only with low [substrates] (*top*) are indicated by the *red arrows*. (***D***) Overlay of the ferryl fractions at different reaction times (*red and blue points*) with the *ΔA*_318_ kinetic traces from stopped-flow experiments under similar reaction conditions (*colored lines*). The ferryl fractions with high [substrates] (*red*) and low [substrates] (*blue*) are determined from the spectra in Figure S9. The blue dotted line is the kinetic trace of the low-substrates condition in the presence of 5% glycerol, which suppresses ferryl reduction to Fe^III^ product(s).

Mössbauer spectra of samples frozen after complete decay of the ferryl complex (**Figure 6C**) reveal the development of broad, paramagnetically split features of high-spin Fe^III^ species (*red arrows*) exclusively under the low-[substrates] conditions (*compare top and bottom spectra*). We further confirmed the formation of high-spin Fe^III^ species by EPR spectroscopy on samples prepared in an identical fashion (**Figure S11**). The *ΔA*318-versus-time traces from stopped-flow experiments carried out under similar reaction conditions (**Figure 6D**) reflect partitioning of the ferryl state between productive decay to transparent Fe^II^ species, which is dominant with high [substrates] (*red trace*), and unproductive decay to UV-absorbing Fe^III^ species, which prevails with low [substrates] (*solid blue trace*).

We serendipitously discovered an additional probe of the non-chlorinating outcome(s) seen at low [substrates], and it provided further evidence that the unproductive pathway involves substrate dissociation. Inclusion of 5% (*v*/*v*) glycerol in the reaction led to more rapid and more nearly complete decay of the UV absorption (**Figure 6D**, *dashed blue trace*), implying enhanced return to Fe^II^ cofactor forms at the expense of Fe^III^ species. In light of our inference that unproductive ferryl decay in the reaction with *d*4-l-Lys is initiated by substrate dissociation (at low [Cl^−^] and [l-Lys]) and suppressed by substrate rebinding (at high [Cl^−^] and [l-Lys]), we posited that this effect of glycerol might reflect its opportunistic reaction with ferryl complex(es) missing one or both substrates. We varied the concentration of the potentially competitive reductant in stopped-flow experiments monitoring either the UV absorption from the ferryl intermediate and ferric product(s) (**Figure 7A**) or the visible absorption from the reactant quinary complex (**Figure 7B**). As expected, the effect depends on the glycerol concentration, with both decay of the ferryl complex and re-development of the quinary complex becoming increasingly rapid with increasing [glycerol]. More tellingly, 1,1,2,3,3-[^2^H5]-glycerol (*d*5-glycerol) is markedly less potent in these effects than the protium-bearing compound (**Figure 7C**, *compare purple and black traces*). This observation implies that the effect of glycerol involves cleavage of one of its C–H bonds, most likely leading to oxidation of one of its three hydroxyl groups to a carbonyl. Transfer of hydrogen to the ferryl complex should require close approach, which should be blocked by the substrates when they are bound. Indeed, glycerol accelerates ferryl decay and suppresses Fe^III^ formation less efficiently at higher [Cl^−^] than at lower [Cl^−^] (**Figure S12**), consistent with the hypothesis that it reacts only with complex(es) lacking the anion. This conclusion suggests the possibility that BesD might, in the presence of low concentrations of *d*4-l-Lys and Cl^−^ to trigger O_2_ activation and ferryl complex formation, be capable of chlorinating non-native substrates that do not efficiently trigger the enzyme. Interestingly, this sort of “bait-and-switch” reactivity was recently demonstrated for BesC, the substrate-triggered, heme-oxygenase-like, nonheme-diiron partner of BesD in the β-ethynyl-l-serine pathway.^36^

**Figure 7.**
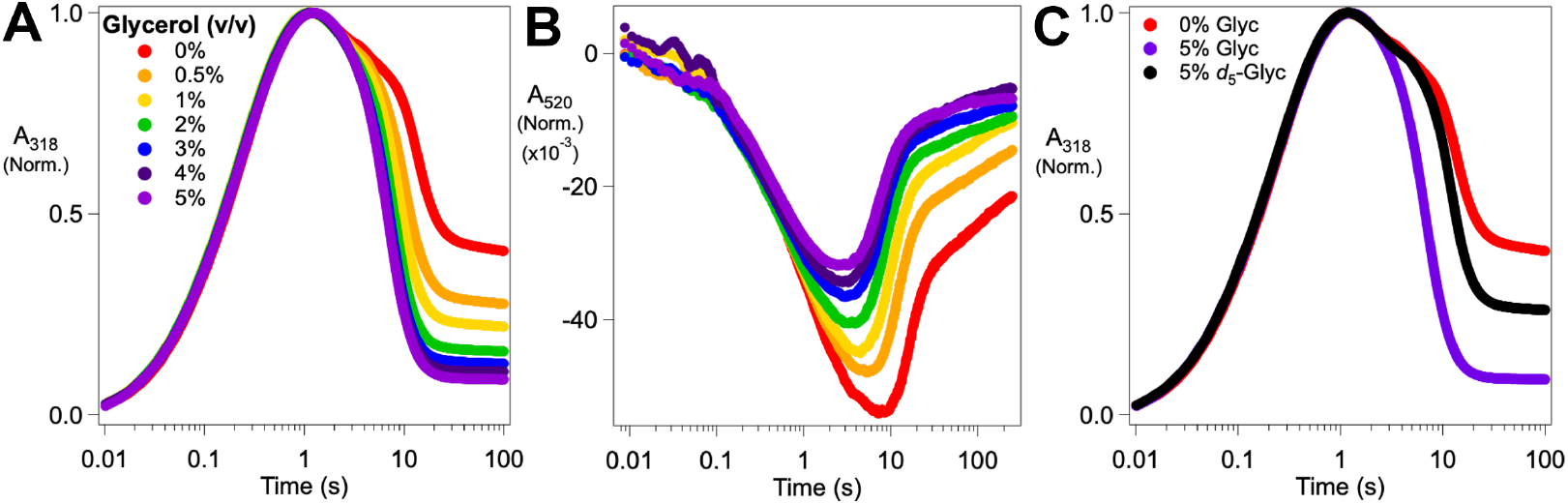
Dependence of the kinetics and outcome of decay of the reactive ferryl state of BesD on glycerol concentration monitored by absorbance (***A***) at 318 nm (ferryl intermediate and ferric products) and (***B***) at 520 nm (ferrous reactant complex). An anoxic solution containing 1.1 mM BesD, 0.8 mM Fe^II^, 10 mM 2OG, 100 mM NaCl, 3 mM *d*_4_-l-Lys, and glycerol (0-5%, *v*/*v*) was mixed at 5 °C with an equal volume of O_2_-saturated buffer (∼ 1.8 mM [O_2_]). Traces in ***A*** and ***B*** were normalized on the basis of the maximum amplitude in ***A***. (***C***) Chloroferryl kinetic traces from ***A*** for 0% glycerol (*red*) and 5% glycerol (*purple*) compared to a sample containing 5% *d*_5_-glycerol (*black*). Raw traces can be found in **Figure S13**

### Cooperative Cl^−^ and l-Lys Binding Also in the Ferryl-mimicking Vanadyl Complex

The vanadyl ion [(V^IV^≡O)^2+^] has emerged over the past several years as a stable structural mimic of the reactive ferryl species in Fe/2OG enzymes.^34,37–39^ It has an *S* = 1/2 ground state that gives rise to an EPR spectrum that is centered at *g* ∼ 2 and dominated by strong and highly anisotropic ^51^V (*I* = 7/2) hyperfine coupling. Binding of vanadyl in an enzyme active site prevents its dimerization to an EPR-inactive state, as it is prone to do in solution. Thus, it can effectively serve as a “turn-on” EPR probe for iron enzymes. In the case of BesD, we found that the positions of its hyperfine features in continuous wave (CW) EPR spectra are sensitive to substrate binding (**Figures 8A** and **S14**). The effect of substrate binding is most evident in the low- and high-field regions (**Figure 8A**, *insets*), reflecting primarily a ∼ 20 MHz diminution of *A*z. However, in the absence of added chloride, 25 mM l-Lys shifts the features of only a small fraction of the total signal from the BesD•[(V^IV^≡O)^2+^]•succinate complex (**Figure 8B**), implying fractional saturation even at this relatively high concentration. This weak binding of the prime substrate would seem to mirror the trends seen for the ferryl intermediate, and, indeed, l-Lys binding to the BesD•(V^IV^≡O)^2+^•succinate complex also exhibits the linkage with chloride binding (**Figure S15**). The observation of this cooperative binding effect for l-Lys and Cl^−^ also in the BesD complex with vanadyl is more anecdotal evidence of its utility as a ferryl mimic. Further studies of the vanadyl probe using advanced pulsed-EPR and protein crystallography may thus afford insight into the prime substrate/anion cooperativity in the chloroferryl state, how substrate positioning is altered by the transition from ferrous reactant to ferryl intermediate state (similar to VioC), and the effect of different halides/pseudohalides on substrate positioning and reactivity.

**Figure 8.**
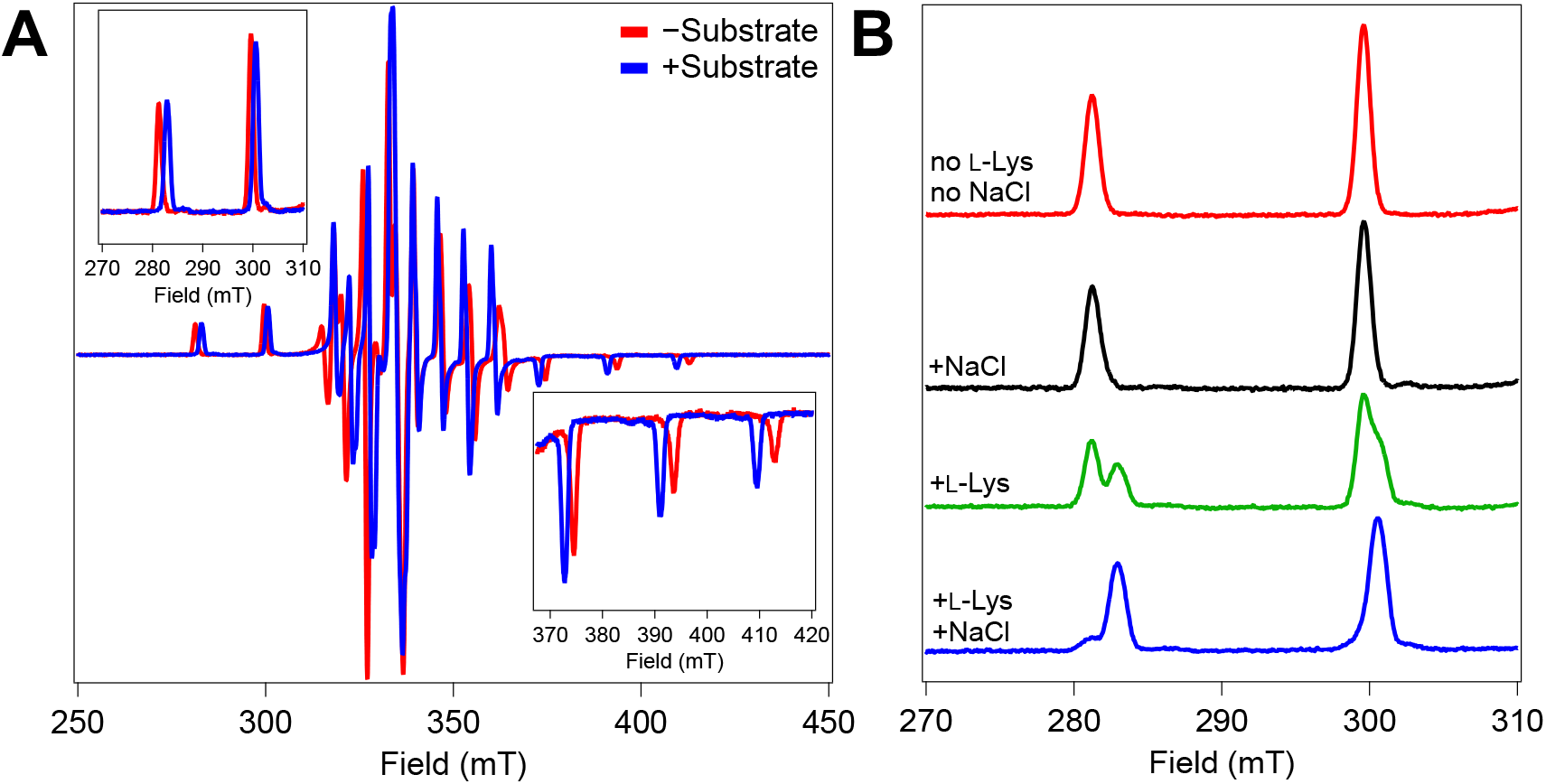
EPR Spectra of BesD•vanadyl•succinate complex with and without substrate present. (*Insets*) Low field and high field portions of the g_z_ component signal. (B) Low field components of BesD•vanadyl•succinate complex in the absence or presence of 25 mM l-Lys and/or 1.0 M NaCl. Experimental EPR parameters are denoted in the *Materials and Methods*.

### Insight into the Orders of Substrate Binding and Product Release; Challenges for Deployment in Biocatalysis

As noted, the Fe/2OG halogenases are prime targets for biocatalysis, because they install an exceptional leaving group that could enable late-stage coupling of advanced synthetic intermediates. In this context, commercial viability will be favored by high total turnover numbers (TON) and turnover frequencies (TOF). An understanding of their chemical and kinetic mechanisms could facilitate optimization of reaction conditions for these key metrics. In the case of BesD, the diminishing halogenation efficiency at low substrate concentrations, which results from the lability of the ferryl intermediate state, would suggest that high [Cl^−^] would be preferable. However, results of LC-MS analysis of reactions carried out under these conditions hinted that, as [Cl^−^] is increased into the molar regime, the quantities of (i) l-Lys consumed, (ii) l-Lys-derived products formed, and (iii) succinate co-product formed are actually modestly diminished (**Figure S16A, B**). These observations implicate substrate (Cl^−^) inhibition in BesD, a phenomenon previously observed in the hydroxylase TauD. In that case, inhibition by the prime substrate, taurine, was attributed to its out-of-sequence binding to the TauD•Fe^II^•succinate complex to delay release of the co-product. We tested for the hallmark of this type of inhibition, the capacity of a substrate to delay rebinding of 2OG (detected by the MLCT band at ∼ 500 nm) at the end of a turnover, for the case of l-Lys and Cl^−^ in the BesD reaction.

Indeed, we found that increasing [Cl^−^] markedly delays re-binding of 2OG, such that, at 1.5 M, the re-development phase of the *ΔA*520-versus-time trace is incomplete after 15 min (**Figure 9A**). l-Lys does not have such an inhibitory effect (**Figure 9B**), despite the apparently general binding synergy between the two substrates. How might this dichotomy arise? As shown in **Figure 2**, although *prior* binding of Cl^−^ and l-Lys to BesD markedly impedes 2OG addition (*red trace*), the presence of the other two substrates does not delay 2OG binding when the enzyme is challenged with all three substrates simultaneously (*purple trace*). Thus, inhibition by Cl^−^ at the end of a turnover must arise from its kinetic trapping of a complex with 4-Cl-l-Lys, succinate, or both still bound in the active site (neglecting the C1-derived CO2 and O_2_-derived water co-products, which are likely to dissociate rapidly). The trapping by Cl^−^ of a complex with 4-Cl-l-Lys bound can plausibly rationalize the failure of l-Lys similarly to inhibit 2OG re-binding: sharing a binding site with 4-Cl-l-Lys, the prime substrate cannot re-bind until the product has dissociated. As long as succinate dissociates rapidly with respect to 4-Cl-l-Lys (either before or after), the prime substrate will not inhibit 2OG re-acquisition, even at high concentration. This conclusion would suggest that, in the reversible association of substrates with Fe^II^ and Fe^IV^ enzyme forms, Cl^−^ leaves before and binds after the prime substrate.

**Figure 9.**
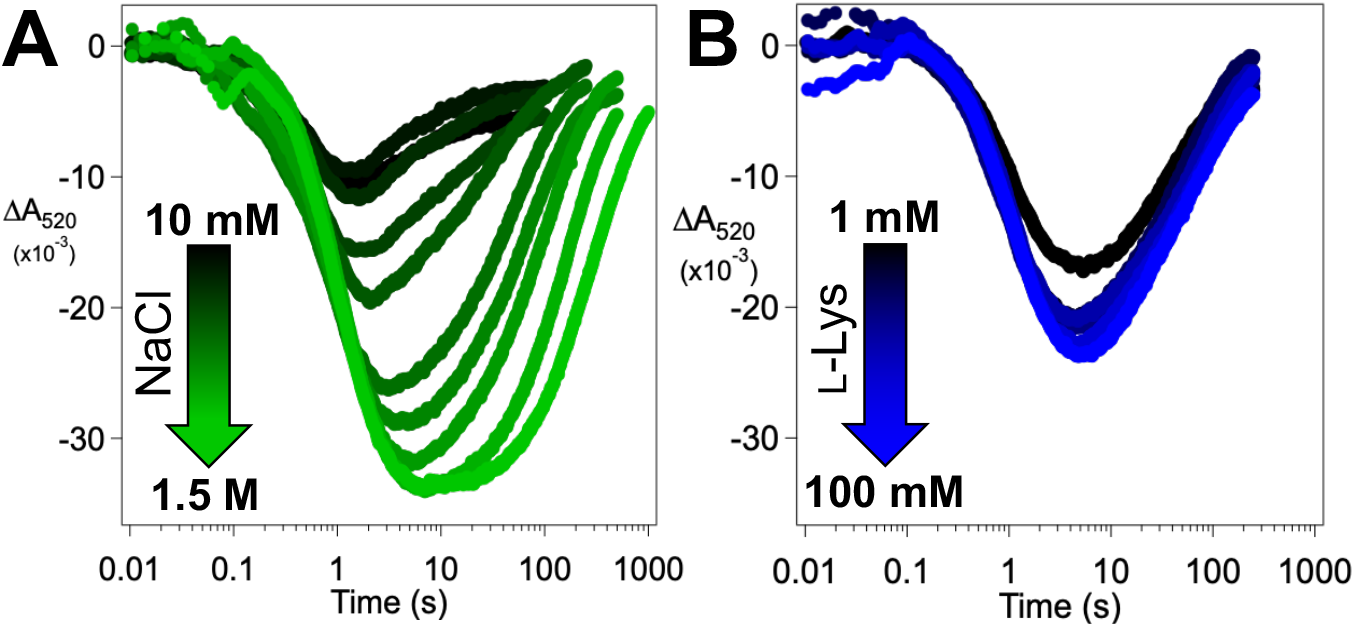
(A) Absorbance-vs-time traces at 520 nm as an anoxic solution of BesD (0.55 mM), Fe^II^ (0.4 mM), 2OG (5 mM), l-Lys (10 mM), and NaCl varying from 20 mM - 3.0 M was mixed at 5°C with an equal volume of air-saturated buffer (∼0.18 mM final [O_2_]). (B) Kinetic traces at 520 nm of similar anoxic solutions as (A), but with a constant 400 mM [NaCl] and varying [l-Lys] from 2 - 200 mM are mixed with air-saturated buffer.

## Conclusions

Investigation of the reaction dynamics of the l-lysine 4-chlorinase BesD has revealed strong heterotropic cooperativity in binding of the prime substrate and the iron-coordinating chloride (or a non-native anion). Unexpectedly, these substrates can dissociate even in the reactive ferryl intermediate state. Thus, when it is challenged with deuterium at the target C–H bond, BesD extensively samples Fe^IV^ forms missing substrates. These forms can either rebind the substrates or decay through pathways leading to univalent reduction to Fe^III^ species or oxidation of a buffer component (glycerol). Because cooperativity is markedly diminished in the ferryl state relative to that in the reactant ferrous state, BesD is prone to uncoupled reactions, in which the ferryl intermediate forms from the O_2_-reactive BesD•Fe^II^•2OG•Cl^−^•l-Lys quinary complex but decays through pathways initiated by Cl^−^ and/or l-Lys dissociation rather than hydrogen abstraction from l-Lys. Efficient chlorination of the deuterium-bearing prime substrate thus requires high concentrations of l-Lys and Cl^−^, but the latter substrate can severely inhibit subsequent turnovers by delaying re-acquisition of 2OG. This understanding lays out challenges and opportunities associated with use of BesD in biocatalysis and criteria to use in comparing it to homologs in searches for more favorable starting points.

## Acknowledgments

This work was supported by the National Institutes of Health (GM138580 to J.M.B. Jr., GM127079 to C.K., GM134271 to M.C.Y.C., GM141284 to A.S., GM119707 to A.K.B.). J.W.S acknowledges support of the National Institute of General Medical Sciences of the National Institute of Health (F32GM136156). M.E.N. acknowledges the support of a National Science Foundation Graduate Research Fellowship. C.-Y.L. was supported by a grant from The Jane Coffin Childs Memorial Fund for Medical Research. The content is solely the responsibility of the authors and does not necessarily represent the official views of the National Institute of Health.

## References

(1) Kovaleva, E. G.; Lipscomb, J. D. Versatility of Biological Non-Heme Fe(II) Centers in Oxygen Activation Reactions. Nat. Chem. Biol. 2008, 4 (3), 186–193. https://doi.org/10.1038/nchembio.71.

(2) Li, F.; Zhang, X.; Renata, H. Enzymatic C–H Functionalizations for Natural Product Synthesis. Curr. Opin. Chem. Biol. 2019, 49, 25–32. https://doi.org/10.1016/j.cbpa.2018.09.004.

(3) Zhang, R. K.; Huang, X.; Arnold, F. H. Selective CH Bond Functionalization with Engineered Heme Proteins: New Tools to Generate Complexity. Curr. Opin. Chem. Biol. 2019, 49, 67– 75. https://doi.org/10.1016/j.cbpa.2018.10.004.

(4) Sheldon, R. A.; Brady, D.; Bode, M. L. The Hitchhiker’s Guide to Biocatalysis: Recent Advances in the Use of Enzymes in Organic Synthesis. Chem. Sci. 2020, 11 (10), 2587– 2605. https://doi.org/10.1039/C9SC05746C.

(5) Solomon, E. I.; Light, K. M.; Liu, L. V.; Srnec, M.; Wong, S. D. Geometric and Electronic Structure Contributions to Function in Non-Heme Iron Enzymes. Acc. Chem. Res. 2013, 46 (11), 2725–2739. https://doi.org/10.1021/ar400149m.

(6) Solomon, E. I.; Goudarzi, S.; Sutherlin, K. D. O2 Activation by Non-Heme Iron Enzymes. Biochemistry 2016, 55 (46), 6363–6374. https://doi.org/10.1021/acs.biochem.6b00635.

(7) Hausinger, R. P. FeII/Alpha-Ketoglutarate-Dependent Hydroxylases and Related Enzymes. Crit. Rev. Biochem. Mol. Biol. 2004, 39 (1), 21–68. https://doi.org/10.1080/10409230490440541.

(8) Dunham, N. P.; Mitchell, A. J.; Del Río Pantoja, J. M.; Krebs, C.; Bollinger, J. M.; Boal, A. K. α-Amine Desaturation of d-Arginine by the Iron(II)- and 2-(Oxo)Glutarate-Dependent l-Arginine 3-Hydroxylase, VioC. Biochemistry 2018, 57 (46), 6479–6488. https://doi.org/10.1021/acs.biochem.8b00901.

(9) Hashimoto, T.; Kohno, J.; Yamada, Y. Epoxidation in Vivo of Hyoscyamine to Scopolamine Does Not Involve a Dehydration Step. Plant Physiol. 1987, 84 (1), 144–147.

(10) Chang, W.; Guo, Y.; Wang, C.; Butch, S. E.; Rosenzweig, A. C.; Boal, A. K.; Krebs, C.; Bollinger, J. M. Mechanism of the C5 Stereoinversion Reaction in the Biosynthesis of Carbapenem Antibiotics. Science 2014, 343 (6175), 1140–1144.

(11) Busby, R. W.; Townsend, C. A. A Single Monomeric Iron Center in Clavaminate Synthase Catalyzes Three Nonsuccessive Oxidative Transformations. Bioorg. Med. Chem. 1996, 4 (7), 1059–1064.

(12) Vaillancourt, F. H.; Bolin, J. T.; Eltis, L. D. The Ins and Outs of Ring-Cleaving Dioxygenases. Crit. Rev. Biochem. Mol. Biol. 2006, 41 (4), 241–267. https://doi.org/10.1080/10409230600817422.

(13) Vaillancourt, F. H.; Vosburg, D. A.; Walsh, C. T. Dichlorination and Bromination of a Threonyl-S-Carrier Protein by the Non-Heme Fe(II) Halogenase SyrB2. Chembiochem Eur. J. Chem. Biol. 2006, 7 (5), 748–752. https://doi.org/10.1002/cbic.200500480.

(14) Mitchell, A. J.; Zhu, Q.; Maggiolo, A. O.; Ananth, N. R.; Hillwig, M. L.; Liu, X.; Boal, A. K. Structural Basis for Halogenation by Iron- and 2-Oxo-Glutarate-Dependent Enzyme WelO5. Nat. Chem. Biol. 2016, 12 (8), 636–640. https://doi.org/10.1038/nchembio.2112.

(15) Matthews, M. L.; Krest, C. M.; Barr, E. W.; Vaillancourt, F. H.; Walsh, C. T.; Green, M. T.; Krebs, C.; Bollinger, J. M. Substrate-Triggered Formation and Remarkable Stability of the C-H Bond-Cleaving Chloroferryl Intermediate in the Aliphatic Halogenase, SyrB2. Biochemistry 2009, 48 (20), 4331–4343. https://doi.org/10.1021/bi900109z.

(16) Neugebauer, M. E.; Sumida, K. H.; Pelton, J. G.; McMurry, J. L.; Marchand, J. A.; Chang, M. C. Y. A Family of Radical Halogenases for the Engineering of Amino-Acid-Based Products. Nat. Chem. Biol. 2019, 15 (10), 1009–1016. https://doi.org/10.1038/s41589-019-0355-x.

(17) Price, J. C.; Barr, E. W.; Tirupati, B.; Bollinger, J. M.; Krebs, C. The First Direct Characterization of a High-Valent Iron Intermediate in the Reaction of an α-Ketoglutarate-Dependent Dioxygenase: A High-Spin Fe(IV) Complex in Taurine/α-Ketoglutarate Dioxygenase (TauD) from Escherichia Coli †. Biochemistry 2003, 42 (24), 7497–7508. https://doi.org/10.1021/bi030011f.

(18) Price, J. C.; Barr, E. W.; Glass, T. E.; Krebs, C.; Bollinger, J. M. Evidence for Hydrogen Abstraction from C1 of Taurine by the High-Spin Fe(IV) Intermediate Detected during Oxygen Activation by Taurine:Alpha-Ketoglutarate Dioxygenase (TauD). J. Am. Chem. Soc. 2003, 125 (43), 13008–13009. https://doi.org/10.1021/ja037400h.

(19) Price, J. C.; Barr, E. W.; Hoffart, L. M.; Krebs, C.; Bollinger, J. M. Kinetic Dissection of the Catalytic Mechanism of Taurine:α-Ketoglutarate Dioxygenase (TauD) from Escherichia Coli. Biochemistry 2005, 44 (22), 8138–8147. https://doi.org/10.1021/bi050227c.

(20) Huang, X.; Groves, J. T. Beyond Ferryl-Mediated Hydroxylation: 40 Years of the Rebound Mechanism and C-H Activation. J. Biol. Inorg. Chem. JBIC Publ. Soc. Biol. Inorg. Chem. 2017, 22 (2–3), 185–207. https://doi.org/10.1007/s00775-016-1414-3.

(21) Blasiak, L. C.; Vaillancourt, F. H.; Walsh, C. T.; Drennan, C. L. Crystal Structure of the Non-Haem Iron Halogenase SyrB2 in Syringomycin Biosynthesis. Nature 2006, 440 (7082), 368. https://doi.org/10.1038/nature04544.

(22) Matthews, M. L. Protein Control of Dioxygen Activation, Substrate-Hydrogen Abstraction, and Ligand-Radical-Transfer Outcome in the Aliphatic Halogenase, SyrB2, Pennsylvania State University, 2011. https://etda.libraries.psu.edu/catalog/12079 (accessed 2019-11-07).

(23) Martinie, R. J.; Livada, J.; Chang, W.; Green, M. T.; Krebs, C.; Bollinger, J. M.; Silakov, A. Experimental Correlation of Substrate Position with Reaction Outcome in the Aliphatic Halogenase, SyrB2. J. Am. Chem. Soc. 2015, 137 (21), 6912–6919. https://doi.org/10.1021/jacs.5b03370.

(24) Ye, S.; Neese, F. Quantum Chemical Studies of C–H Activation Reactions by High-Valent Nonheme Iron Centers. Curr. Opin. Chem. Biol. 2009, 13 (1), 89–98. https://doi.org/10.1016/j.cbpa.2009.02.007.

(25) Matthews, M. L.; Neumann, C. S.; Miles, L. A.; Grove, T. L.; Booker, S. J.; Krebs, C.; Walsh, C. T.; Bollinger, J. M. Substrate Positioning Controls the Partition between Halogenation and Hydroxylation in the Aliphatic Halogenase, SyrB2. Proc. Natl. Acad. Sci. 2009, 106 (42), 17723–17728. https://doi.org/10.1073/pnas.0909649106.

(26) Hillwig, M. L.; Liu, X. A New Family of Iron-Dependent Halogenases Acts on Freestanding Substrates. Nat. Chem. Biol. 2014, 10 (11), 921–923. https://doi.org/10.1038/nchembio.1625.

(27) Marchand, J. A.; Neugebauer, M. E.; Ing, M. C.; Lin, C.-I.; Pelton, J. G.; Chang, M. C. Y. Discovery of a Pathway for Terminal-Alkyne Amino Acid Biosynthesis. Nature 2019, 567 (7748), 420–424. https://doi.org/10.1038/s41586-019-1020-y.

(28) Matthews, M. L.; Chang, W.; Layne, A. P.; Miles, L. A.; Krebs, C.; Bollinger Jr, J. M. Direct Nitration and Azidation of Aliphatic Carbons by an Iron-Dependent Halogenase. Nat. Chem. Biol. 2014, 10 (3), 209–215. https://doi.org/10.1038/nchembio.1438.

(29) Yang, J.; Yang, Z.; Yin, Y.; Rao, M.; Liang, Y.; Ge, M. Three Novel Polyene Macrolides Isolated from Cultures of Streptomyces Lavenduligriseus. J. Antibiot. (Tokyo) 2016, 69 (1), 62–65. https://doi.org/10.1038/ja.2015.76.

(30) Kissman, E. N.; Neugebauer, M. E.; Sumida, K. H.; Swenson, C. V.; Sambold, N. A.; Marchand, J. A.; Millar, D. C.; Chang, M. C. Y. Biocatalytic Control of Site-Selectivity and Chain Length-Selectivity in Radical Amino Acid Halogenases. Proc. Natl. Acad. Sci. 2023, 120 (12), e2214512120. https://doi.org/10.1073/pnas.2214512120.

(31) Pratter, S. M.; Light, K. M.; Solomon, E. I.; Straganz, G. D. The Role of Chloride in the Mechanism of O2 Activation at the Mononuclear Nonheme Fe(II) Center of the Halogenase HctB. J. Am. Chem. Soc. 2014, 136 (26), 9385–9395. https://doi.org/10.1021/ja503179m.

(32) Munck, E. Chapter 6: Aspects of 57Fe Mössbauer Spectroscopy. In Physical Methods in Bioinorganic Chemistry: Spectroscopy and Magnetism; Que Jr., L., Ed.; University Science, 2000.

(33) Krebs, C.; Galonić Fujimori, D.; Walsh, C. T.; Bollinger, J. M. Jr. Non-Heme Fe(IV)–Oxo Intermediates. Acc. Chem. Res. 2007, 40 (7), 484–492. https://doi.org/10.1021/ar700066p.

(34) Dunham, N. P.; Chang, W.-C.; Mitchell, A. J.; Martinie, R. J.; Zhang, B.; Bergman, J. A.; Rajakovich, L. J.; Wang, B.; Silakov, A.; Krebs, C.; Boal, A. K.; Bollinger, J. M. Two Distinct Mechanisms for C-C Desaturation by Iron(II)- and 2-(Oxo)Glutarate-Dependent Oxygenases: Importance of α-Heteroatom Assistance. J. Am. Chem. Soc. 2018, 140 (23), 7116–7126. https://doi.org/10.1021/jacs.8b01933.

(35) Pan, J.; Wenger, E. S.; Matthews, M. L.; Pollock, C. J.; Bhardwaj, M.; Kim, A. J.; Allen, B. D.; Grossman, R. B.; Krebs, C.; Bollinger, J. M. Evidence for Modulation of Oxygen Rebound Rate in Control of Outcome by Iron(II)- and 2-Oxoglutarate-Dependent Oxygenases. J. Am. Chem. Soc. 2019, 141 (38), 15153–15165. https://doi.org/10.1021/jacs.9b06689.

(36) McBride, M. J.; Nair, M. A.; Sil, D.; Slater, J. W.; Neugebauer, M. E.; Chang, M. C. Y.; Boal, A. K.; Krebs, C.; Bollinger, J. M. Substrate-Triggered μ-Peroxodiiron(III) Intermediate in the 4-Chloro-l-Lysine-Fragmenting Heme-Oxygenase-like Diiron Oxidase (HDO) BesC: Substrate Dissociation from, and C4 Targeting by, the Intermediate. Biochemistry 2022, 61 (8), 689–702. https://doi.org/10.1021/acs.biochem.1c00774.

(37) Martinie, R. J.; Pollock, C. J.; Matthews, M. L.; Bollinger, J. M.; Krebs, C.; Silakov, A. Vanadyl as a Stable Structural Mimic of Reactive Ferryl Intermediates in Mononuclear Nonheme-Iron Enzymes. Inorg. Chem. 2017, 56 (21), 13382–13389. https://doi.org/10.1021/acs.inorgchem.7b02113.

(38) Davis, K. M.; Altmyer, M.; Martinie, R. J.; Schaperdoth, I.; Krebs, C.; Bollinger, J. M.; Boal, A. K. Structure of a Ferryl Mimic in the Archetypal Iron(II)- and 2-(Oxo)-Glutarate-Dependent Dioxygenase, TauD. Biochemistry 2019, 58 (41), 4218–4223. https://doi.org/10.1021/acs.biochem.9b00598.

(39) Mitchell, A. J.; Dunham, N. P.; Martinie, R. J.; Bergman, J. A.; Pollock, C. J.; Hu, K.; Allen, B. D.; Chang, W.-C.; Silakov, A.; Bollinger, J. M.; Krebs, C.; Boal, A. K. Visualizing the Reaction Cycle in an Iron(II)- and 2-(Oxo)-Glutarate-Dependent Hydroxylase. J. Am. Chem. Soc. 2017, 139 (39), 13830–13836. https://doi.org/10.1021/jacs.7b07374.

