## Supplemental Information for "Synergistic Binding of the Halide and Cationic Prime Substrate of the l-Lysine 4-Chlorinase, BesD, in Both Ferrous and Ferryl States"

### Materials and Methods

Materials

All materials were obtained from Sigma-Aldrich, Fisher Scientific, or VWR unless otherwise noted. All solutions were prepared using 18.2 MΩ/cm deionized water (Barnstead).

Cloning, Expression, and Purification of BesD

The gene for *Streptomyces lavenduligriseus* BesD was excised via a double-restriction digest, NdeI/XhoI (New England Biolabs), from a pET16-His-PrecissionCutSite-IMPDH expression vector and ligated (T4 Ligase, NEB) into a pET26b expression vector with a stop codon preceding the vector-based C-terminal His-tag. This BesD construct lacking affinity tags, displayed below with start/stop codons underlined, was verified by DNA sequencing at the Molecular Core Facility of The Pennsylvania State University.

ATGAGCAGCAATGGGCAGGAAAGCACCGTCGTCAATCCTCTGGAGCAGGGCGCACTGCG
CCGTATGGCGCATCACTACCACCGGTACGGCATCGCCACCGTCACCGATCTGATTCGGGA
AGACGTCCGCAAAAACGTGCGGGCGGAGGCGGACCGGCTGCTGGAGAAATACGCCGAGC
GGCGTGATCTCCGCCTCCAGACCACCGGATACACCCGCCGCTCGATGTCCGTGGTGCAG
AGCGAGACGATCGCGGGCAACAGCGAGCTGGTCACCTCGATCTACGCGAATCCGGAACT
GCTCGGCGCGCTGGAGCGCATAGCCGGCGAGAAACTGCACCCCTGCCCCAAGGCCGAC
GAGGAATTCCTCATCACCCGGCAGGAACGGAGCGGGGACACGCACGGCTGGCACTGGGG
CGATTTCAGCTTCGCCCTCATCTGGGTGCTCCAGGCCCCGCCCATCGACATCGGCGGCAT
GCTCCAGTGCGTACCGCACACGGAGTGGGACAAGTCCGATCCGCGGATCCACCAGTATCT
CGTCGACAATCCCATCCACACGTACCACTTCCAGTCGGGCGACGTGTATTTCCTGCGCAC
CGACACCACGCTCCACCGCACGGTCCCGCTGCGCGAGGACACCACCCGCATCATCCTGA
ACATGACCTGGGCGGGCGAGCGGGACCTGCGGCGCGAGCTCAAGGGCGACGACCGCTG
GTGGGAGGACGCCGACGTGCCGGCAGCCGCTGCACTGGATGACTGA

The plasmid containing the tag-less BesD construct was introduced via heat-shock transformation into BL21 (DE3) *E. coli* cells (New England Biolabs). A single colony was used to inoculate a starter culture of rich Luria-Bertani (LB) growth medium supplemented with 70 μg/mL kanamycin antibiotic and left to grow for 16-18 hours at 37°C and 200 RPM. Large culture flasks containing rich-LB and kanamycin were inoculated with 10 mL of starter culture per 1 L growth medium and grown at 37°C and 200 RPM until an $OD_{600}$ ~ 1.2-1.5 was reached. Flasks were cooled on ice for 30 minutes before BesD overexpression was initiated by the addition of IPTG to a final concentration of 0.25 mM. Cultures were left to overexpress overnight at 18°C and 200 RPM for 16-18 hours. Cells were harvested by centrifugation at 5,000 xg and flash frozen in $LN_2$ before storage at -80°C.

Purification of BesD was carried out between 4-10°C. Frozen cell paste was thawed and resuspended in 5 mL/g lysis buffer containing 100 mM HEPES, pH 7.5. A spatula tip (~0.1 g) each of lysozyme and DNase I was added to the thawed cell mixture. Resuspended cells were lysed by sonification at 50% amplitude at 10 s/30 s on/off intervals for seven minutes. The cell lysate was then diluted by 50% with a saturated ammonium sulfate ($AmSO_4$) solution. The lysate was incubated for one hour at 4°C with stirring before centrifugation at 22,000 xg for 35 minutes. Due to the extreme solubility of BesD in high salinity solutions, BesD remained in the soluble supernatant and cellular debris/salt-precipitated proteins was pelleted. The supernatant was decanted and diluted again by 50% using 100 mM HEPES, pH 7.5 buffer to lower the $AmSO_4$ to a final concentration of 25% saturation. This step was necessary before concentration as 10 kDa molecular weight cutoff (MWCO) concentrators (Merck Millipore Ltd) run extremely slow at >25%

$AmSO_4$. The lysate supernatant was concentrated to a manageable volume before overnight dialysis in 100 mM HEPES, pH 7.5 to remove the remaining $AmSO_4$.

The desalinated lysate was concentrated after dialysis to <100 mL and loaded onto a self-packed Q column (26/100, 600 mL resin) for ion exchange purification with an AKTA Purifier FPLC system (GE Healthcare). The protein was eluted using a gradient from 0-50% buffer A (100 mM HEPES, pH 7.5, 50 mM $MgSO_4$) to buffer B (100 mM HEPES, 1 M $MgSO_4$, pH 7.5) with a final wash of 100% buffer B. Fractions containing BesD were pooled and concentrated via centrifugal concentrators. Purified BesD was dialyzed against a solution of 5 mM EDTA and 100 mM HEPES, pH 7.5 buffer for 4+ hours and two additional dialysis steps without EDTA to remove exogenous metals. Protein purity was accessed by SDS-PAGE (as shown in Figure S17) and protein concentration was determined by its absorbance at 280 nm using a calculated molar absorptivity of 50,420 $M^{-1}$ $cm^{-1}$ (https://web.expasy.org/protparam/). Typical protein yields ~250 mg BesD per liter of rich LB medium.

Anion Exchange Chromatography to Remove Chloride Ions from Lysine Stocks

To better control chloride concentrations under various conditions, it was necessary to remove the chloride ions from commercial lysine stocks. While protiated L-lysine (Sigma-Aldrich) was available as a free amino acid, i.e. chloride-free, no lysine deuterated isotopes could be found without HCl counterions.

The $d_4$-4,4,5,5-L-Lys•2HCl stock (Cambridge Isotopes) was dissolved in a 25:75 mixture of 1M NaOH:100 mM HEPES, pH 7.5 buffer to a final concentration of 0.5 M $d_4$-L-Lys, pH 6-8. Final pH was confirmed to be neutral via pH strips (EMD Millipore). A 5-mL FF Q-column (Cytiva) was prepared by rinsing the column with 25 mL of 1 M ammonium acetate, pH 6.5, followed by 25 mL of $H_2O$ via syringe or peristaltic pump. The high excess of the weaker-binding acetate ions outcompeted and exchanged with the resin-bound chloride ions.[1] The $d_4$-L-Lys stock was loaded onto and washed off the column with 10 mL $H_2O$ and the flowthrough was collected. Chloride ions in the L-lysine solution were expected to bind to the Q resin and release the previously resin-bound acetate ions. The flowthrough solution was flash frozen in $LN_2$ and lyophilized for >18 hours. The solid was resuspended in $H_2O$ as $d_4$-4,4,5,5-L-Lys •2NaAcetate. Removal of chloride ions was confirmed by addition of a high concentration of treated lysine stock to BesD-$Fe^{II}$-2OG complex in a UV-Vis spectrophotometer, as shown in Figure S18. Lack of perturbation of the $Fe^{II}$-2OG MLCT at 520 nm indicates little to no chloride ions present in the lysine stock.

Absorption Spectroscopy

UV-Visible spectra were measured using an Agilent 8453 UV-visible spectroscopy system housed in an MBraun (Stratham, NH) anoxic chamber. Titrations of the various analytes were carried out using two identical cuvettes; one for the sample of interest and a control lacking $Fe^{II}$ for subtraction during the analysis. Spectra were collected after each addition of an aliquot of the concentrated stock titrant solution. Each experimental spectrum was corrected for dilution by the titrant stock solution and the initial spectrum ([titrant]= 0 M) was subtracted from the dilution-corrected spectrum. The absorbance at 800 nm was subtracted from each spectrum and set to zero and, if required, were baseline corrected via a point-based cubic spline in Kazan Viewer.[2]

Microscale Thermophoresis

Affinity measurements for free L-lysine binding to BesD was carried out using a NanoTemper Tech. Monolith NT.115 microscale thermophoresis (MST) instrument. A stock of BesD was labelled with a fluorescent moiety via coupling to protein-based amines using the protein labeling kit RED-NHS 2nd generation (NanoTemper Tech.). MST samples were prepared in a anaerobic chamber. The fluorophore labelled BesD (RED-BesD) was supplemented to the protein binding mixture. The protein solution was aliquoted and added to an L-Lys concentration series in a 1:5 mixture for a final concentration of 50 nM RED-BesD, 0.5 mM BesD, 0.5 mM $Fe^{II}$, 2.5 mM 2OG,

0.05% TWEEN, and 100 mM HEPES, pH 7.5. L-lysine concentration ranges varied based on the concentration of chloride present in each series to better incapsulate the binding range. Unlabeled BesD was added to the samples to prevent unincorporated ferrous ions from causing fluctuations in the fluorescence signal due to quenching. Samples were loaded into capillary tubes and sealed with wax on both ends to prevent sample oxidation. Samples were analyzed by MST using 100% MST power and 100% excitation power at 650 nm and emission at 670 nm.

Stopped-Flow Absorption Spectroscopy

Stopped-flow absorption experiments were performed on an Applied Photophysics Ltd. (Leatherhead, UK) SX-20 stopped-flow spectrophotometer housed inside a glovebox. Reactions were carried out at 5°C in a single-mixing configuration with a 1 cm (or 0.2 cm, for reactions in **Figure 6D**) pathlength and photomultiplier tube (PMT) detector. Wavelengths were selected/isolated from the broadband light source before the reaction cell via a monochromator. Specific experimental details are provided in the figure legends.

LC-MS Activity Assays as a Function of Chloride Concentration

Reactions to assess the activity of BesD under single- and multi-turnover conditions were carried out. Multi-turnover assays contained 1 mM substrate (L-Lys or *d*$_4$-L-Lys), 0.1 mM $(NH_4)_2Fe(SO_4)_2{\bullet}6H_2O$, 0.1 mM BesD, 5 mM $Na_2$2OG, 5 mM ascorbic acid, and NaCl at varying concentrations. Reactions were initiated by addition of 2OG, mixed thoroughly to allow for proper oxygenation in ambient air, and allowed to proceed for 20 min at room temperature before quenching with a 2-fold dilution in methanol. After quenching, internal standards of the opposite substrate isotopologue (L-Lys for *d*$_4$-L-Lys reactions, *d*$_4$-L-Lys for L-Lys reactions) and sodium *d*$_4$-succinate was added to the mixture before filtration. The samples were immediately transferred and analyzed by LS-MS, as described below.

Single turnover assays contained 1 mM substrate (L-Lys or *d*$_4$-L-Lys), 0.5 mM $(NH_4)_2Fe(SO_4)_2{\bullet}6H_2O$, 0.7 mM BesD, 0.375 mM $Na_2$2OG, and NaCl at varying concentrations. Reactions were prepared inside an anaerobic chamber and initiated by the addition of $O_2$-saturated buffer (5°C) for a final concentration of 0.6 mM. The reactions were allowed to proceed for 5 min before quenching with a 2-fold dilution in methanol. After quenching, internal standards of the opposite substrate isotopologue (L-Lys for *d*$_4$-L-Lys reactions, *d*$_4$-L-Lys for L-Lys reactions) and sodium *d*$_4$-succinate was added to the mixture before filtration. The samples were immediately transferred and analyzed by LS-MS, as described below.

Liquid chromatography coupled to mass spectrometry (LC-MS) analysis was carried out on a 1200 series LC system connected to a 6400 series triple quadrupole mass spectrometer (Agilent Technologies). Analysis of L-Lys-based products by LC-MS was performed by injection (5 μL) of reaction mixture onto a SeQuant ZIC-HILIC (3.5 μM, 150 x 2.1 mm; EMD Millipore) column with a mobile phase consisting of 10:90 (v/v) MeCN:$H_2O$, 10 mM $NH_4HCO_2$, pH 5.0 (A) and 90:10 (v/v) MeCN:$H_2O$, 10 mM $NH_4HCO_2$, pH 5.0 (B). Separation of elutants was carried out with a linear gradient of 95% B to 30% B over 25 min followed by a linear gradient of 30% B to 95% B over 5 min at a flow rate of 0.2 mL/min. Samples were analyzed in positive-ionization mode.

Evaluation of succinate and 2OG present in reactions was performed by injection (2 μL) onto an Extend-C18 (1.8 μM, 50 x 4.6 mm; Agilent) column with a mobile phase consisting of 0.1% formic acid in $H_2O$ (A) and MeCN (B). An isocratic method of 5% B at 0.3 mL/min was utilized to separate elutants over a 5 min period. Samples were analyzed in negative-ionization mode.

Freeze-Quench Mössbauer Spectroscopy

Freeze-quench Mössbauer samples were prepared according to previously published procedures.[3] Mössbauer spectra were recorded on a spectrometer from SEECO (Edina, MN)

equipped with a Janis SVT-400 variable-temperature cryostat. The reported isomer shift is given relative to the centroid of the spectrum of α-iron metal at room temperature. External magnetic fields were applied parallel to the direction of propagation of the γ-radiation. Simulations of the Mössbauer spectra were carried out using WMOSS spectral analysis software from SEECO (www.wmoss.org, SEE Co., Edina, MN).

Samples were generated from anoxic solutions of 1.8 mM BesD, 1.5 mM $^{57}Fe(II)$, 16 mM 2OG, 10 mM or 60 mM $d_4$-L-Lys, and 100 mM or 3.0 M NaCl in a butter solution of 100 mM HEPES, pH 7.5. These samples were mixed at 5 °C with an equal volume of the buffer that had been saturated with $O_2$ (~1.8 mM). This reaction mixture was allowed to incubate for the varying reaction times indicated in **Figure 6** and subsequently frozen by injection into cryogenically cooled (-150 °C) 2-methylbutane (for reaction times <30 seconds) or by pipetting into a Mössbauer cell cooled on a metal block that was in contact with $LN_2$ (for reaction times >30 seconds). Hand-quench EPR samples were made in parallel under same conditions as Mössbauer samples. Samples were hand-mixed and transferred to EPR tubes before freezing in $LN_2$ at appropriate quench times.

Electron Paramagnetic Resonance Spectroscopy

EPR samples were transferred into custom X-band EPR tubes (Quartz Scientific, Inc., Fairport Harbor, Ohio), and continuous-wave X-band EPR spectra were recorded on a Magnettech MS5000X spectrometer equipped with an Oxford Instruments ESR-900 continuous flow cryostat and an Oxford Instruments ITC-300 temperature controller. Data were collected for vanadyl-based samples at 35 K from 250 to 450 mT and iron-based samples were collected at 10 K from 50 to 425 mT. Spectra were collected with modulation amplitude of 1 mT, microwave power of 0.1 mW, and a microwave frequency of 9.43516 GHz.

BesD-vanadyl samples were prepared with 0.5 mM vanadyl sulfate, 0.6 mM BesD, 10 mM sodium succinate, and 0.4 M sucrose (cryosolvent) in buffer containing 100 mM HEPES, pH 7.5. Concentrations of L-Lys and NaCl for samples are denoted in figures. Analysis of the L-Lys/NaCl titration data was performed using three reference spectra multiplied by a fractionation coefficient in a linear regression to determine the amount of each species in the spectrum. This was performed in the software package Igor Pro 9 (WaveMetrics, Inc., Lake Oswego, OR, USA.)

The collected spectra were baseline corrected by subtraction of a fitted polynomial in Kazan Viewer.[2] The baseline-corrected EPR spectra were simulated with the EasySpin toolbox[4] within MATLAB.

Analysis of L-Lysine and Chloride Cooperativity in $Fe^{II}$-BesD-2OG Complex

As the affinity of L-lysine (L-Lys) and chloride to $Fe^{II}$-BesD-2OG are independently weak but relatively stronger in the presence of the opposite cosubstrate, it is assumed that they bind in a positively cooperative manner. To model this, a square scheme has been constructed as demonstrated below:

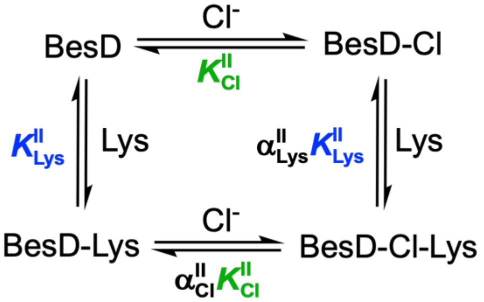


2OG and $Fe^{II}$ are present for all species but omitted for simplicity. The dissociation equilibrium constants, $K_D$, are represented for each substate as either $K^{II}_{Lys}$ or $K^{II}_{Cl}$. The constants for the

second binding events, $Fe^{II}$-Cl + Lys and $Fe^{II}$-Lys + Cl, are described by the first binding event constants, $K^{II}_{Lys}$ and $K^{II}_{Cl}$, multiplied by the cooperativity coefficient, $\alpha^{II}_{Lys}$ and $\alpha^{II}_{Cl}$, respectively. The combined binding events create a thermodynamic cycle that can be represented as:

$$K^{II}_{Cl}\,\alpha^{II}_{Lys}\,K^{II}_{Lys} = K^{II}_{Lys}\,\alpha^{II}_{Cl}\,K^{II}_{Cl}$$

Therefore, the two cooperativity coefficients are identical and can be simplified to a single cooperativity coefficient, $\boldsymbol{\alpha}^{\mathbf{II}}$.

$$\alpha^{II}_{Lys} = \alpha^{II}_{Cl} \quad \equiv \quad \boldsymbol{\alpha}^{\mathbf{II}}$$

This value should be <<1 for strong, positive cooperativity (as observed for chloride binding in the absence/presence of L-lysine in **Figure 1**). Due to the thermodynamic cycle, only three parameters ($K^{II}_{Cl}$, $K^{II}_{Lys}$, and $\boldsymbol{\alpha}^{\mathbf{II}}$) are required to characterize the equilibria among the relevant species. While the first binding events of just L-lysine or just chloride can be determined by a simple binding curve in the absence of the other cosubstate, the parameters of the synergistic second binding event require a more complicated analysis. For example, the titration curve of chloride in the presence of a static concentration of L-lysine, reveals the apparent $K_D$ for chloride binding, $K^{II}_{Cl,app}$, by monitoring the metal-to-ligand charge-transfer (MLCT) band at 520 nm exhibited specifically by a chloride ion bound to a $Fe^{II}$-BesD-2OG complex (i.e., BesD-Cl and BesD-Cl-Lys, from the above scheme). This constant can be expressed as:

$$K^{II}_{Cl,app} = \frac{([BesD]+[BesD\cdot Lys])[Cl]}{([BesD\cdot Cl]+[BesD\cdot Lys\cdot Cl])}$$

Using the definitions of $K^{II}_{Lys}$, $K^{II}_{Cl}$, and $\boldsymbol{\alpha}^{\mathbf{II}}K^{II}_{Cl}K^{II}_{Lys}$, this expression can be substituted and rearranged:

$$K^{II}_{Cl,app} = \frac{\left([BesD]+[BesD]\frac{[Lys]}{K^{II}_{Lys}}\right)[Cl]}{\left(\frac{[BesD][Cl]}{K^{II}_{Cl}}+\frac{[BesD][Lys][Cl]}{\boldsymbol{\alpha}^{\mathbf{II}}K^{II}_{Cl}K^{II}_{Lys}}\right)} = \frac{\left(1+\frac{[Lys]}{K^{II}_{Lys}}\right)[BesD][Cl]}{\left(\frac{1}{K^{II}_{Cl}}+\frac{[Lys]}{\boldsymbol{\alpha}^{\mathbf{II}}K^{II}_{Cl}K^{II}_{Lys}}\right)[BesD][Cl]} = \frac{\left(1+\frac{[Lys]}{K^{II}_{Lys}}\right)K^{II}_{Cl}}{\left(1+\frac{[Lys]}{\boldsymbol{\alpha}^{\mathbf{II}}K^{II}_{Lys}}\right)}$$

Since L-Lys is a weak and strong binder in the absence and presence of Cl-, we assume that $[Lys] \ll K^{II}_{Lys} \ll \frac{[Lys]}{\boldsymbol{\alpha}^{\mathbf{II}}}$, such that $\frac{[Lys]}{K^{II}_{Lys}} \ll 1$ and $\frac{[Lys]}{\boldsymbol{\alpha}^{\mathbf{II}}K^{II}_{Lys}} \gg 1$:

$$\Rightarrow \quad K^{II}_{Cl,app} \approx \frac{K^{II}_{Cl}}{\frac{[Lys]}{\boldsymbol{\alpha}^{\mathbf{II}}K^{II}_{Lys}}} = K^{II}_{Cl}\frac{\boldsymbol{\alpha}^{\mathbf{II}}K^{II}_{Lys}}{[Lys]}$$

This expression allows for the determination of the cooperative binding step, $\boldsymbol{\alpha}^{\mathbf{II}}K^{II}_{Lys}$, by measuring $K^{II}_{Cl,app}$ with varying [Lys] (as observed in **Figure 1**), and using $K^{II}_{Cl}$.

Analysis of productive ferryl intermediate formation and decay

The $\Delta A_{318}$-vs-time traces reporting the formation and decay of the ferryl intermediate in the reaction of BesD with protium substrate were analyzed according to the equation and associated kinetic model shown below. The quinary complex (A) reacts to form the ferryl intermediate (B) and subsequently decays into the product state (C), irreversibly.

$$\mathrm{A} \xrightarrow{k_1} \mathrm{B} \xrightarrow{k_2} \mathrm{C}$$

For the purposes of this analysis, the ferryl intermediate is designated as the only state with a transient absorbance at 318 nm. From this, the following equation can be utilized where $\Delta A$ is the change in amplitude (dictated by concentration and extinction coefficient of the intermediate) and $k_1$ and $k_2$ are the effective formation and decay rate constants, respectively:

$$\Delta A_{318}(\mathrm{t}) = \Delta A\left(\frac{k_1}{k_2 - k_1}\right)\left(e^{-k_1 t} - e^{-k_2 t}\right)$$

Analysis of productive and unproductive ferryl intermediate decay

To extract the formation and decay rates for the ferryl intermediate in BesD when two reaction outcomes are present, a step-wise, branching pathway is invoked, as the ferryl can either decay productively or unproductively. Typically, the ferryl intermediate can be analyzed using an expression derived from a simple A to B to C model (shown above), as only the ferryl intermediate (B) absorbs appreciably at 318 nm. In the case of BesD with deuterated substrates, the ferryl appears to decay via two different pathways: either to a ferrous product (C) or ferric product (D) state. The ferric state has an appreciable absorbance at 318 nm overlapping with the ferryl LMCT and requires a more complex expression to account for the mixed absorbance of ferryl intermediate and ferric product. The model used for this stepwise branching pathway is demonstrated below:

$$\mathrm{A} \xrightarrow{k_1} \mathrm{B} \begin{array}{l} \xrightarrow{k_2} \mathrm{C} \\ \xrightarrow{k_3} \mathrm{D} \end{array}$$

The overall decay of the ferryl intermediate (B) is a combination of both productive and unproductive pathways and is represented by $k_{\mathrm{decay}}$.

$$k_{\mathrm{decay}} = k_2 + k_3$$

The integrated rate laws for the four states are as follows:

$$[\mathrm{A}]_t = [\mathrm{A}]_0\, e^{-k_1 t}$$

$$[\mathrm{B}]_t = \frac{[\mathrm{A}]_0 k_1}{k_{\mathrm{decay}} - k_1}\left(e^{-k_1 t} - e^{-k_{\mathrm{decay}} t}\right)$$

$$[\mathrm{C}]_t = \frac{[\mathrm{A}]_0 k_1 k_2}{k_{\mathrm{decay}} - k_1}\left(\frac{1 - e^{-k_1 t}}{k_1} - \frac{1 - e^{-k_{\mathrm{decay}} t}}{k_{\mathrm{decay}}}\right)$$

$$[\mathrm{D}]_t = \frac{[\mathrm{A}]_0 k_1 k_3}{k_{\mathrm{decay}} - k_1}\left(\frac{1 - e^{-k_1 t}}{k_1} - \frac{1 - e^{-k_{\mathrm{decay}} t}}{k_{\mathrm{decay}}}\right)$$

As stated previously, the absorbance at 318 nm as a function of time is defined as a combination of ferryl (B) and ferric (D) multiplied by their respective extinction coefficients.

$$\Delta A_{318} = \varepsilon_B[B]_t + \varepsilon_D[D]_t$$

Substitution and rearrangement leads to the final analytical expression:

$$\Delta A_{318}(t) = \frac{[A]_0 k_1}{k_{decay} - k_1}\left(\varepsilon_B\left(e^{-k_1 t} - e^{-k_{decay} t}\right) + \varepsilon_D k_3\left(\frac{1 - e^{-k_1 t}}{k_1} - \frac{1 - e^{-k_{decay} t}}{k_{decay}}\right)\right)$$

From this expression and the experimental data measured at $A_{318}$, $k_{decay}$ can be determined for the ferryl intermediate and used for further analysis (see below).

<u>Analysis of $d_4$-L-lysine and NaCl Cooperativity in [$Fe^{IV}$=O]-BesD Complex</u>

The decay rate of the ferryl intermediate in BesD can be affected by both concentration of chloride and $d_4$-L-lysine. At low concentrations of both substrates, the ferryl decays faster and there is a net positive absorbance in the baseline at 318 nm, indicative of the ferryl uncoupling and ending in irreversibly formed ferric state that has an increased absorbance at 318 nm. As [$d_4$-L-Lys] and [$Cl^-$] increase, the ferryl decay rate slows and the $\Delta A_{318}$ in the baseline after ferryl decay is smaller, indicating less uncoupling. Based on these observations and what has been observed in the ferrous state, the following scheme representing the mechanism after oxygen activation by the BesD quinary complex was constructed (with the presence of BesD implicitly inferred):

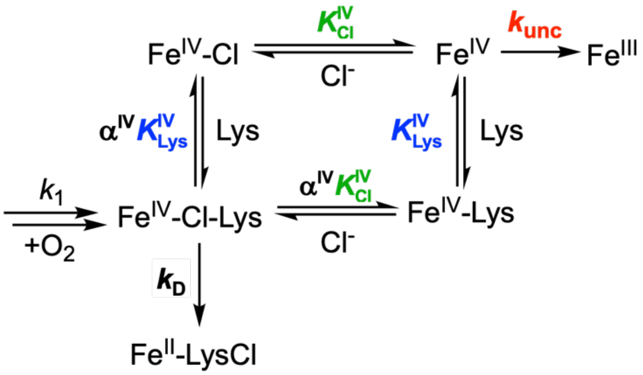


Similar to the ferrous cooperativity scheme, $K^{IV}_{Lys}$ and $K^{IV}_{Cl}$ represent the dissociation constants for L-lysine and chloride, respectively, and $\boldsymbol{\alpha^{IV}}$ is the cooperativity coefficient in the ferryl state. Two rate constants are assigned to the observed two pathways by which the ferryl decays: the rate of uncoupling, $k_{unc}$, and the rate of productive decay, $\boldsymbol{k_D}$. From this scheme, an analytical solution can be derived to express the five parameters that dictate ferryl decay in BesD. As the ferryl absorbance at 318 nm originates from the ligand-to-metal charge transfer (LMCT) band of the oxo ligand and the $Fe^{IV}$, we can assume that each state of the ferryl has a relatively equal extinction coefficient. The experimentally observed $\Delta A_{318}$ is therefore an observation of the total ferryl, as defined below:

$$[Fe^{IV}]_{total} = [Fe^{IV}\text{-Cl-Lys}] + [Fe^{IV}\text{-Lys}] + [Fe^{IV}\text{-Cl}] + [Fe^{IV}]$$

The differential rate law for total ferryl decay can be written as:

$$\frac{d[Fe^{IV}]_{total}}{dt} = -k_{decay}[Fe^{IV}]_{total}$$

The decay of the ferryl only occurs from two of the four ferryl states, $Fe^{IV}$-Cl-Lys, and $Fe^{IV}$. As such, this can be rewritten as:

$$-\frac{d[Fe^{IV}]_{total}}{dt} = -\frac{d[Fe^{IV}]}{dt} - \frac{d[Fe^{IV}\text{-Cl-Lys}]}{dt}$$

$$= k_{unc}[Fe^{IV}] + \boldsymbol{k_D}[Fe^{IV}\text{-Cl-Lys}]$$

If there are fast equilibria for L-Lys and $Cl^-$ binding with respect to the decay rates, the following assumptions can be made,

$$[Fe^{IV}\text{-Cl}] = \frac{[Cl]}{K_{Cl}^{IV}}[Fe^{IV}],$$

$$[Fe^{IV}\text{-Lys}] = \frac{[Lys]}{K_{Lys}^{IV}}[Fe^{IV}],$$

$$[Fe^{IV}\text{-Cl-Lys}] = \frac{[Lys][Cl]}{\alpha^{IV} K_{Lys}^{IV} K_{Cl}^{IV}}[Fe^{IV}]$$

Using the previous definitions, the rate of decay can be rewritten and substituted,

$$k_{unc}[Fe^{IV}] + \boldsymbol{k_D}[Fe^{IV}\text{-Cl-Lys}] = k_{decay}[Fe^{IV}]_{total}$$

$$\Rightarrow \left(k_{unc} + \boldsymbol{k_D}\frac{[Lys][Cl]}{\alpha^{IV} K_{Lys}^{IV} K_{Cl}^{IV}}\right)[Fe^{IV}] = k_{decay}\left(1 + \frac{[Cl]}{K_{Cl}^{IV}} + \frac{[Lys]}{K_{Lys}^{IV}} + \frac{[Lys][Cl]}{\alpha^{IV} K_{Lys}^{IV} K_{Cl}^{IV}}\right)[Fe^{IV}]$$

$$\Rightarrow \quad k_{decay} = \frac{k_{unc} + \boldsymbol{k_D}\frac{[Lys][Cl]}{\alpha^{IV} K_{Lys}^{IV} K_{Cl}^{IV}}}{1 + \frac{[Cl]}{K_{Cl}^{IV}} + \frac{[Lys]}{K_{Lys}^{IV}} + \frac{[Lys][Cl]}{\alpha^{IV} K_{Lys}^{IV} K_{Cl}^{IV}}}$$

From this final analytical expression, the experimentally measured rate of ferryl decay can be fit as a function of [Lys] and [Cl] (as shown in **Figure 5**) to determine the rate of uncoupling, $k_{unc}$, the rate of productive decay, $\boldsymbol{k_D}$, the dissociation constants for L-lysine and chloride, $K_{Lys}^{IV}$ and $K_{Cl}^{IV}$, and the cooperativity coefficient for L-lysine and chloride binding, $\boldsymbol{\alpha^{IV}}$.

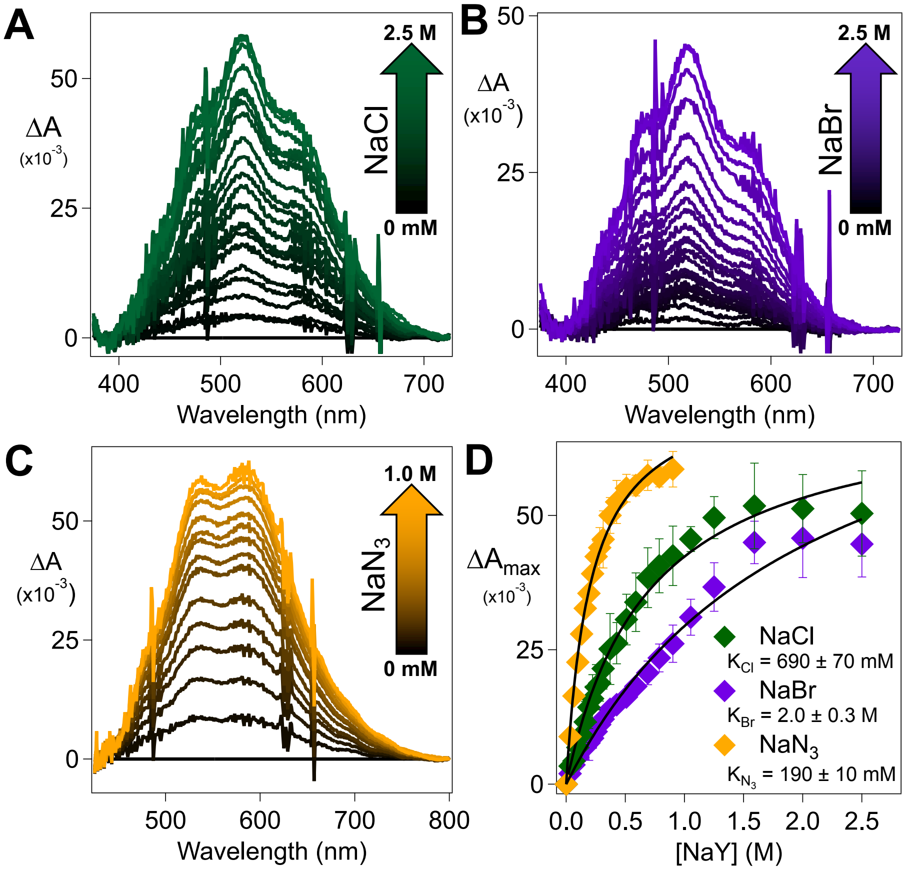


**Figure S1.** (A/B/C) UV-visible absorbance difference spectra of BesD (1.0 mM), $Fe^{II}$ (0.72 mM), and 2OG (5 mM) titrated with a concentrated stock of (A) NaCl, (B) NaBr, or (C) $NaN_3$ in the absence of L-Lys. (D) Titration curve for $A_{520}$ (NaCl), $A_{515}$ (NaBr), and $A_{528}$ ($NaN_3$) as a function of NaY concentration (n=3). $K^{II}_{Cl}$, $K^{II}_{Br}$, and $K^{II}_{N_3}$ were determined by fitting the data to hyperbolic curves.

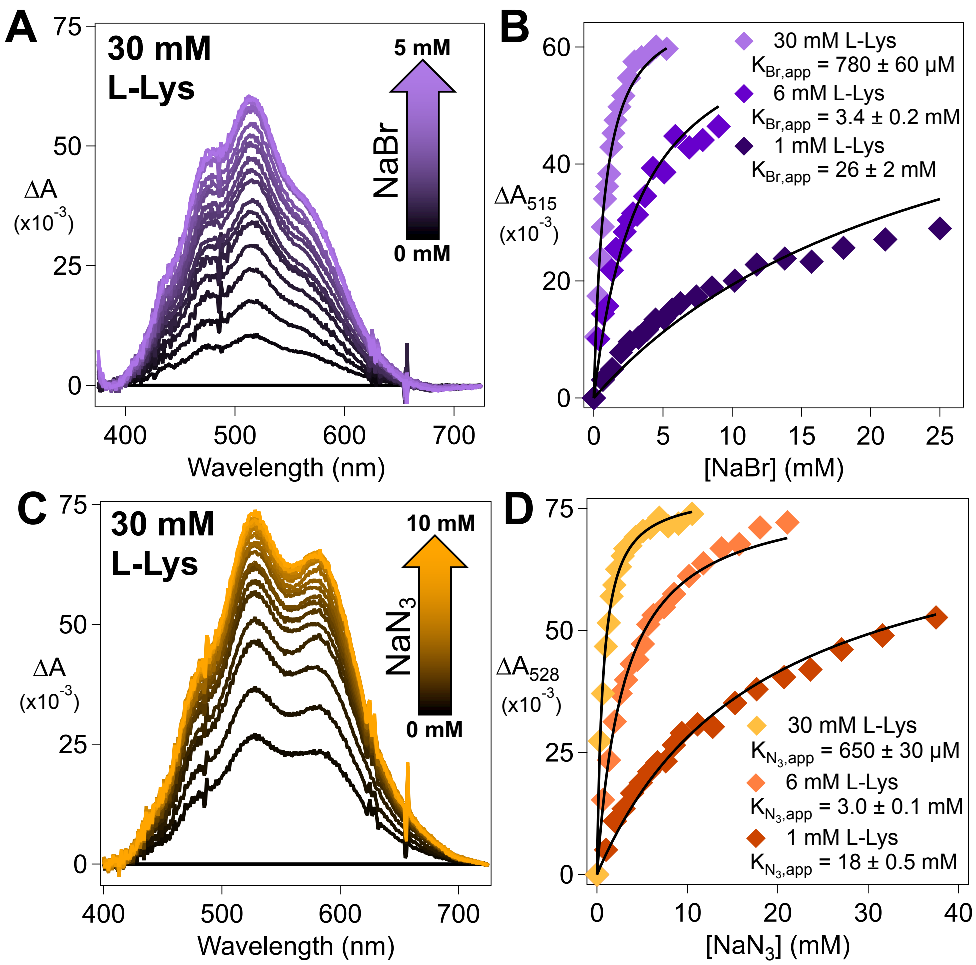


**Figure S2.** (A) UV-visible absorbance difference spectra of BesD (1.0 mM), $Fe^{II}$ (0.72 mM), and 2OG (5 mM) titrated with a concentrated stock of NaBr in the presence of 30 mM L-Lys. (B) Titration curves for $A_{515}$ as a function of NaBr concentration at 1 mM, 6 mM, and 30 mM L-Lys. $K^{II}_{Br,app}$ was determined by fitting the data to a hyperbolic or quadratic curve where appropriate. (C) UV-visible absorbance difference spectra of BesD (1.0 mM), $Fe^{II}$ (0.72 mM), and 2OG (5 mM) titrated with a concentrated stock of NaBr in the presence of 30 mM L-Lys. (D) Titration curves for $A_{528}$ as a function of $NaN_3$ concentration at 1 mM, 6 mM, and 30 mM L-Lys. $K^{II}_{N_3,app}$ was determined by fitting the data to a hyperbolic or quadratic curve where appropriate.

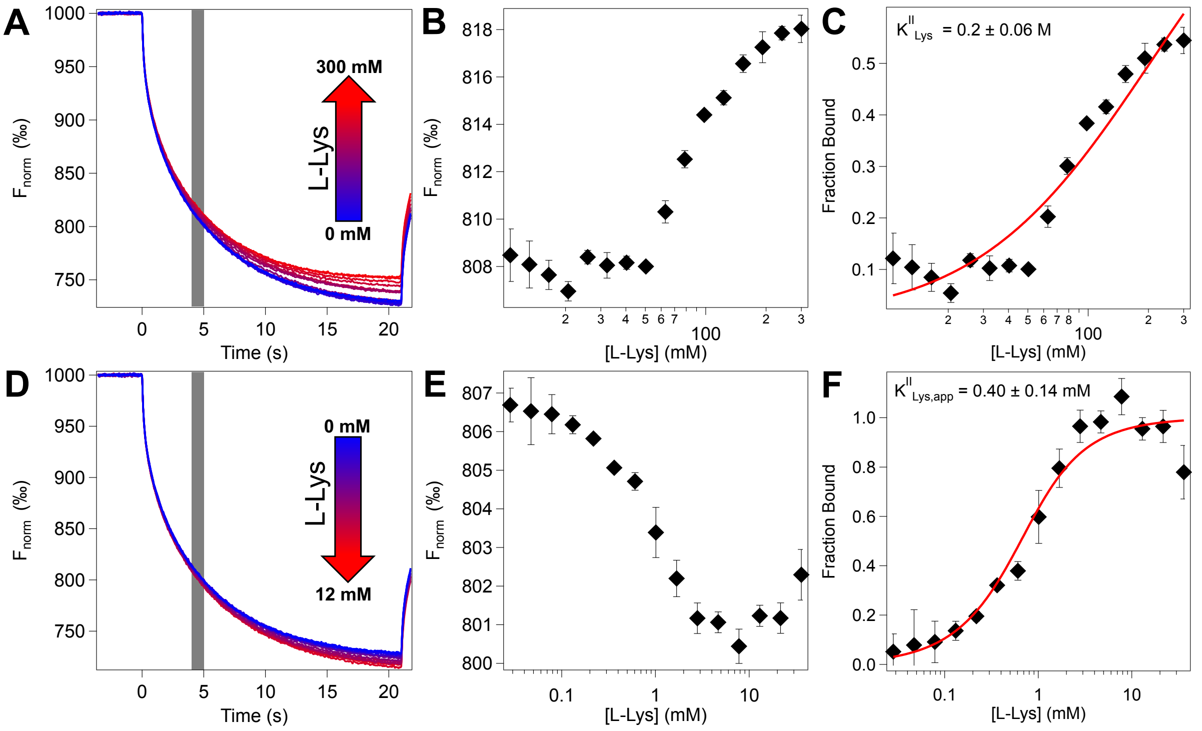


**Figure S3.** (A) MST traces monitoring the thermophoresis of fluorophore-labelled BesD at varying concentrations of L-Lys in the absence of chloride ions. (B) The normalized fluorescence ($F_{Norm}$) between 4-5 seconds of each trace (grey region in A) was extracted and averaged and plotted against [L-Lys] (n=3) (C) Binding curve for L-Lys to BesD in the absence of chloride after iteratively fitting to a hyperbolic curve and conversion to fraction bound based on fit min/max parameters. (D) MST traces monitoring the thermophoresis of fluorophore-labelled BesD at varying concentrations of L-Lys in the presence of 6 mM NaCl. (E) The normalized fluorescence ($F_{Norm}$) between 4-5 seconds of each trace (grey region in D) was extracted and averaged and plotted against [L-Lys] (n=3) (F) Binding curve for L-Lys to BesD in the presence of 6 mM NaCl after iteratively fitting to a quadratic curve and conversion to fraction bound based on fit min/max parameters.

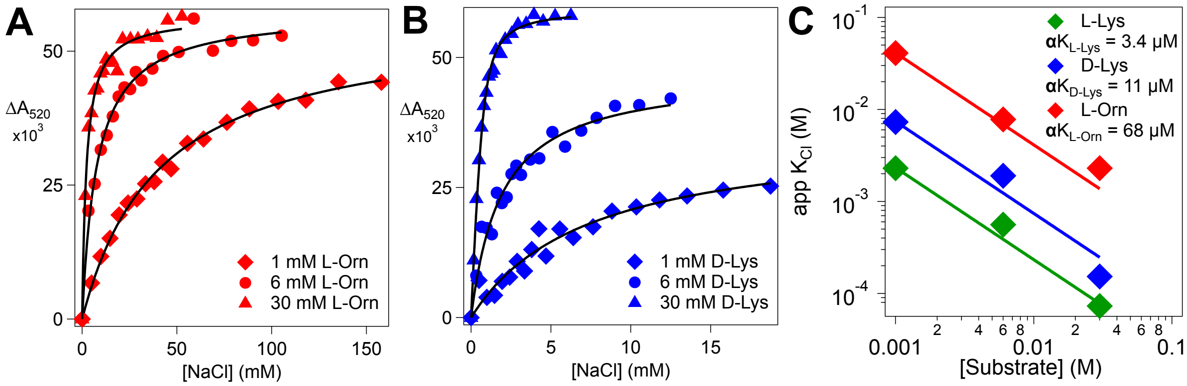


**Figure S4.** Titration curves for $A_{520}$ as a function of [$Cl^-$] at 1 mM, 6 mM, and 30 mM (A) L-Orn and (B) D-Lys. $K_{Cl,app}$ was determined by fitting the data to a hyperbolic or quadratic curve where appropriate. (C) Cooperativity plots for apparent $K_{Cl}$ as a function of total [substrate] for L-Lys (*green*), D-Lys (*blue*), and L-Orn (*red*). The data were fit by equation 1 with constant $K_{Cl}^{II}$ = 0.69 M.

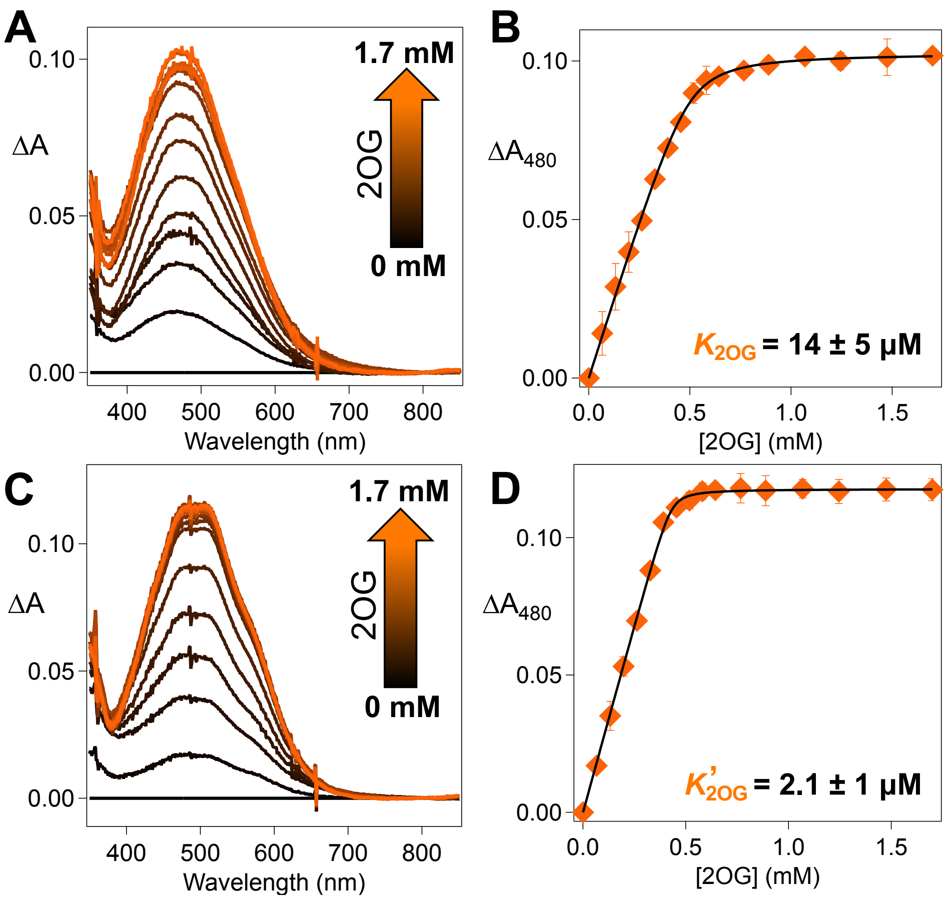


**Figure S5.** (A) UV-visible absorbance difference spectra of BesD (0.5 mM) and $Fe^{II}$ (0.5 mM) titrated with a concentrated stock of 2OG in the absence of L-Lys and $Cl^-$. (B) Titration curves for $A_{480}$ as a function of 2OG concentration. $K_{2OG}$ was determined by fitting the data to a quadratic curve (n=2). (C) UV-visible absorbance difference spectra of BesD (0.5 mM) and $Fe^{II}$ (0.5 mM) titrated with a concentrated stock of 2OG in the presence of 40 mM L-Lys and 1 M $Cl^-$. Each spectrum was acquired after a 30 second incubation period to allow for 2OG to fully bind after each addition (D) Titration curves for $A_{480}$ as a function of 2OG concentration. $K'_{2OG}$ was determined by fitting the data to a quadratic curve (n=3).

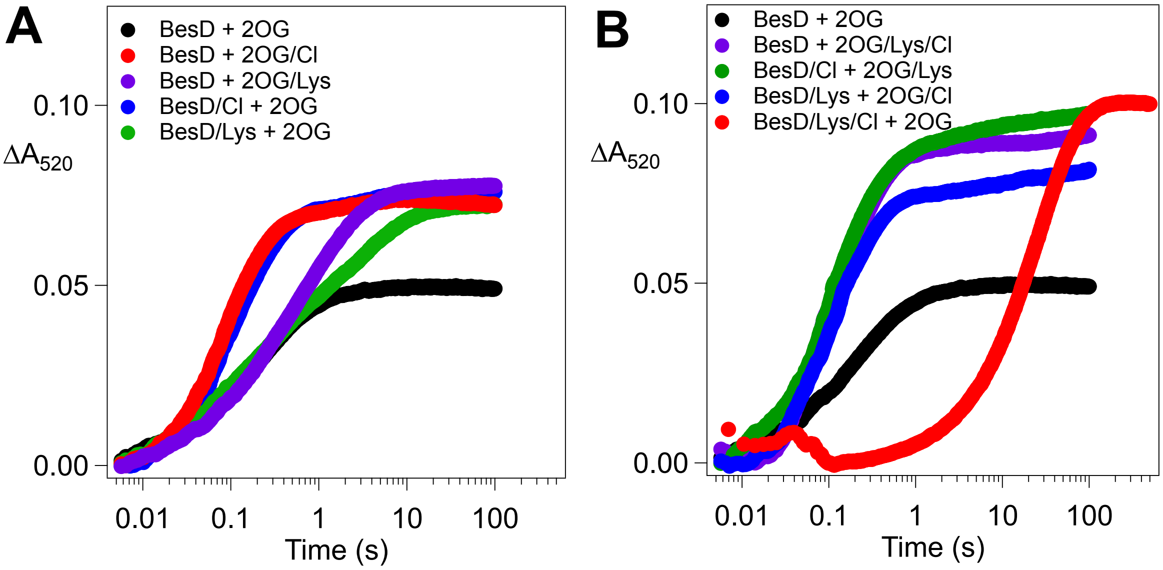


**Figure S6.** Kinetic traces monitoring binding of 2OG (2.4 mM) to BesD (1.1 mM) and $Fe^{II}$ (0.8 mM) at 520 nm. (A) Mixing of BesD with 2OG in the presence of either L-Lys (80 mM) or $Cl^-$ (2 M) by either premixing with BesD or 2OG. (B) Mixing of BesD with 2OG in the presence of both L-Lys and $Cl^-$ in the various mixing permutations with neither, one or the other, or both L-Lys/$Cl^-$ being premixed with BesD or with 2OG. Signals ~30 ms are attributed to mixing artifacts. (B has been recreated from **Figure 2** for comparison)

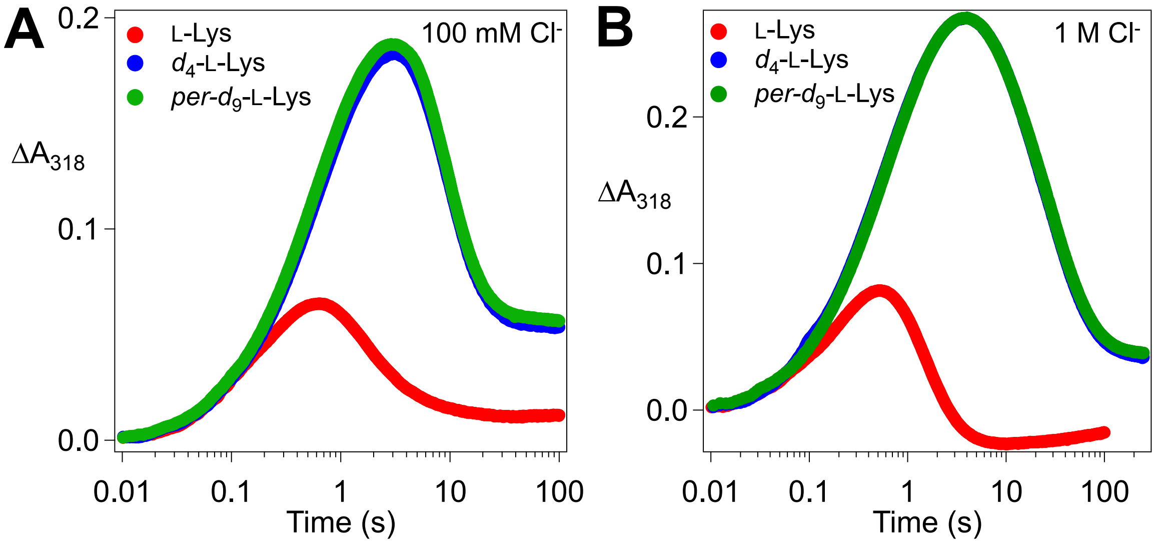


**Figure S7.** (A) Kinetics of the BesD chloroferryl intermediate formation and decay in reactions with L-Lys (red), *d*4-L-Lys (blue), and *per*-*d*9-L-Lys (green) at final concentrations of 100 mM chloride as monitored by its absorbance at 318 nm. (B) Kinetics of the BesD chloroferryl intermediate formation and decay in reactions with L-Lys (red), *d*4-L-Lys (blue), and *per*-*d*9-L-Lys (green) at final concentrations of 1 M chloride as monitored by its absorbance at 318 nm. An anoxic solution of BesD (0.55 mM), $Fe^{II}$ (0.4 mM), 2OG (5 mM), L-Lys/*d*4-L-Lys/ *per*-*d*9-L-Lys (10 mM), and 200 mM NaCl or 2 M NaCl was mixed at 5°C with an equal volume of air-saturated buffer (~0.18 mM final [$O_2$]).

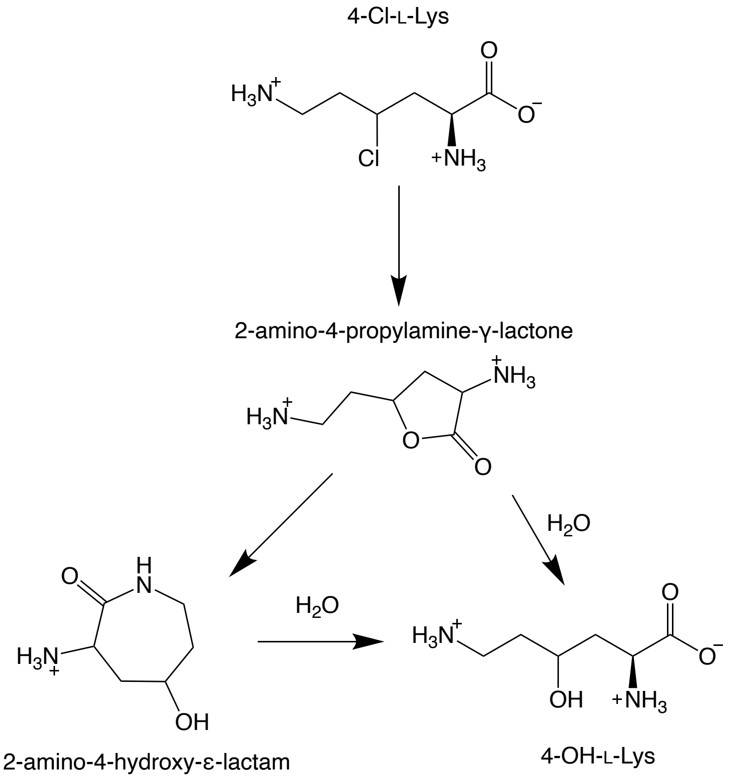


**Figure S8.** Decomposition pathway of 4-Cl-L-Lys into lactam/lactone compounds and finally 4-OH-L-Lys as determined previously.[4]

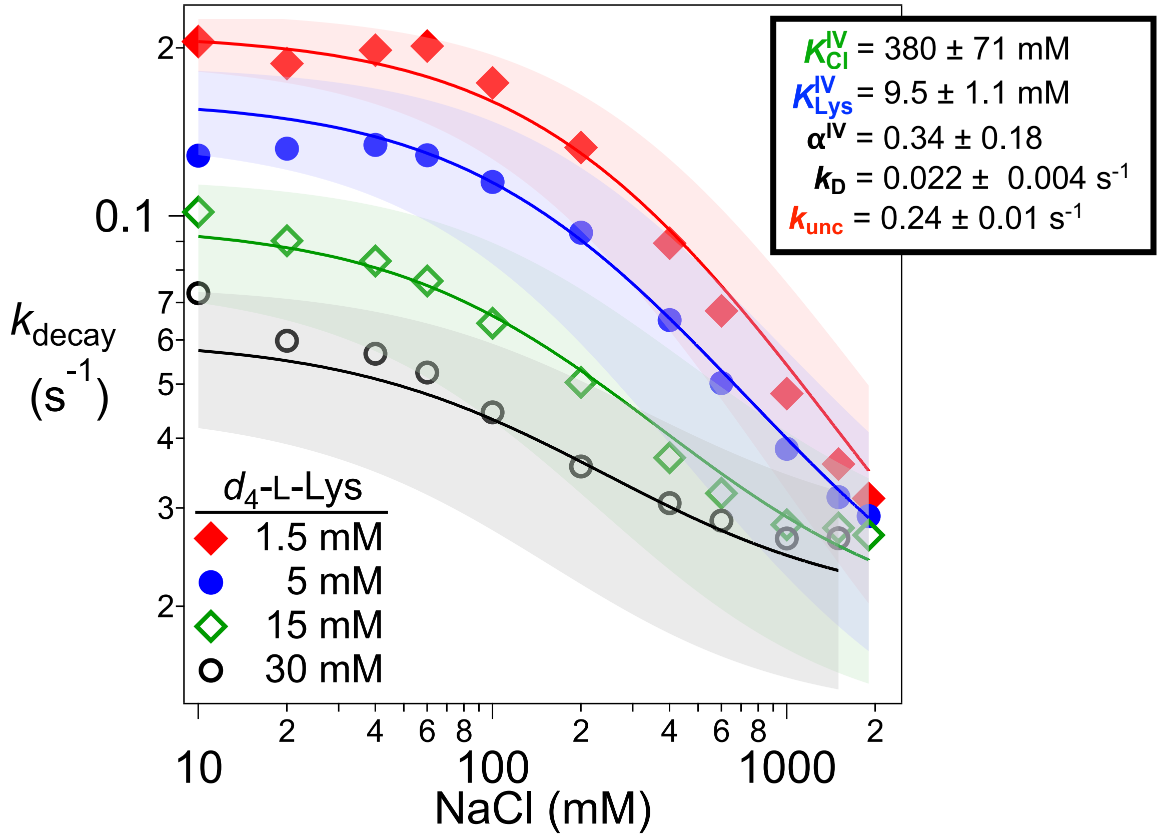


**Figure S9.** Observed decay rates of the ferryl intermediate as a function of [NaCl] at four $d_4$-L-Lys concentrations. Traces were globally fit using equation 2 with the five coefficients linked. Shaded regions represent the uncertainty as the lowest (-) and highest (+) coefficient values determined by the global fit.

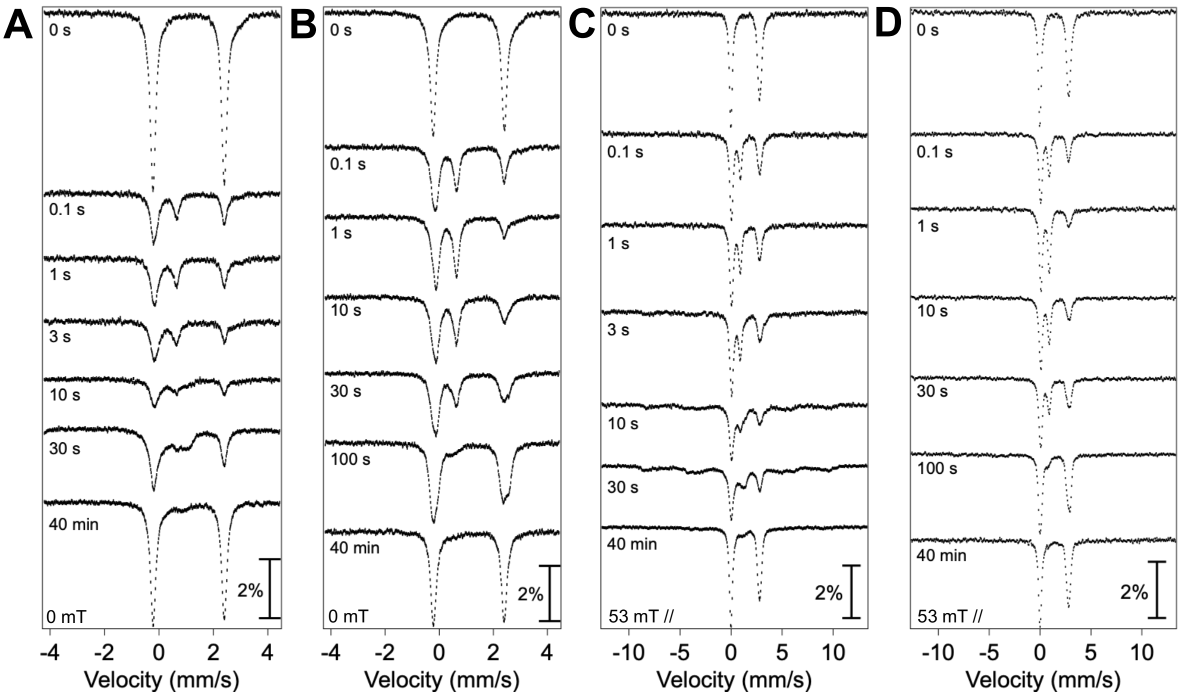


**Figure S10.** Mössbauer spectra (4.2 K) of rapid freeze-quench samples prepared by reacting the BesD quinary complex ($d_4$-L-Lys) at 5°C with $O_2$-saturated buffer. Reaction times are indicated. Samples were prepared with final concentrations of 0.75 mM $^{57}Fe$(II), 0.9 mM BesD, 8 mM 2OG, and (A & C) 5 mM $d_4$-L-Lys and 50 mM NaCl or (B & D) 30 mM $d_4$-L-Lys and 1.5 M NaCl.

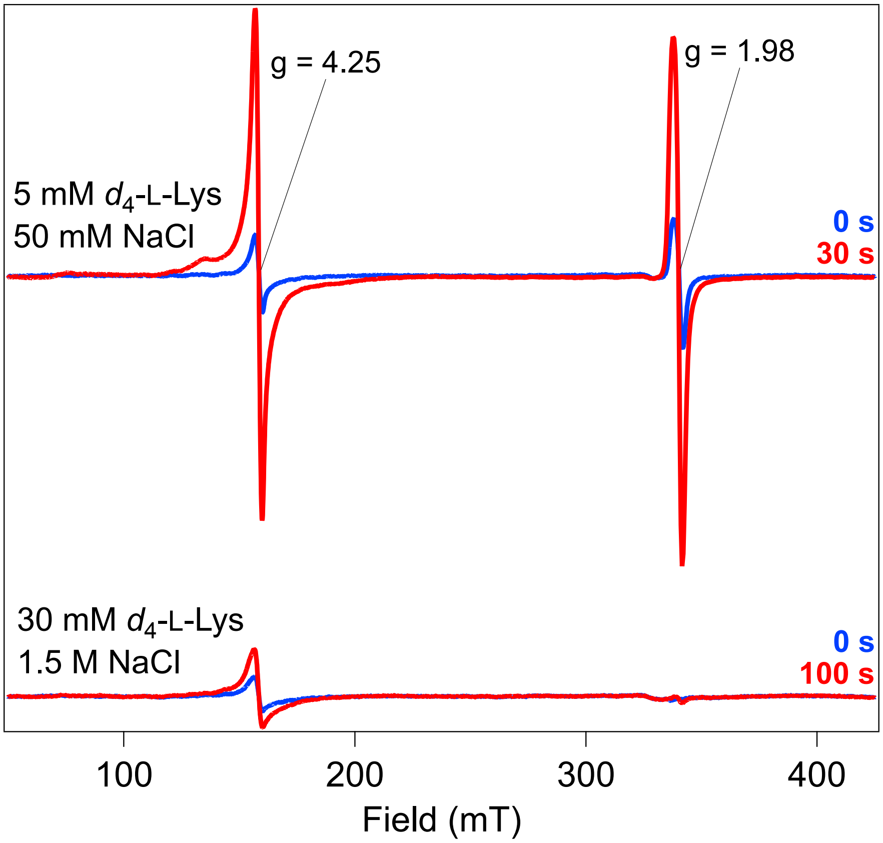


**Figure S11.** Hand-quenched EPR samples made in parallel to Mossbauer samples prepared in **Figure 8**. (*Top*) Reactions under low substrate/anion conditions prepared at 0 s (prepared without $O_2$) and quenched 30 seconds after mixing with saturated-$O_2$ buffer. Initial signal at g = 4.25 and 1.98 in 0 s sample is likely due to $O_2$ contamination during sample preparation. (*Bottom*) Reactions under high substrate/anion conditions prepared at 0 s and 100 s. Spectra were collected at 10 K with 1 mT modulation amplitude, 20 dB microwave power, and a microwave frequency of 9.435165 mW.

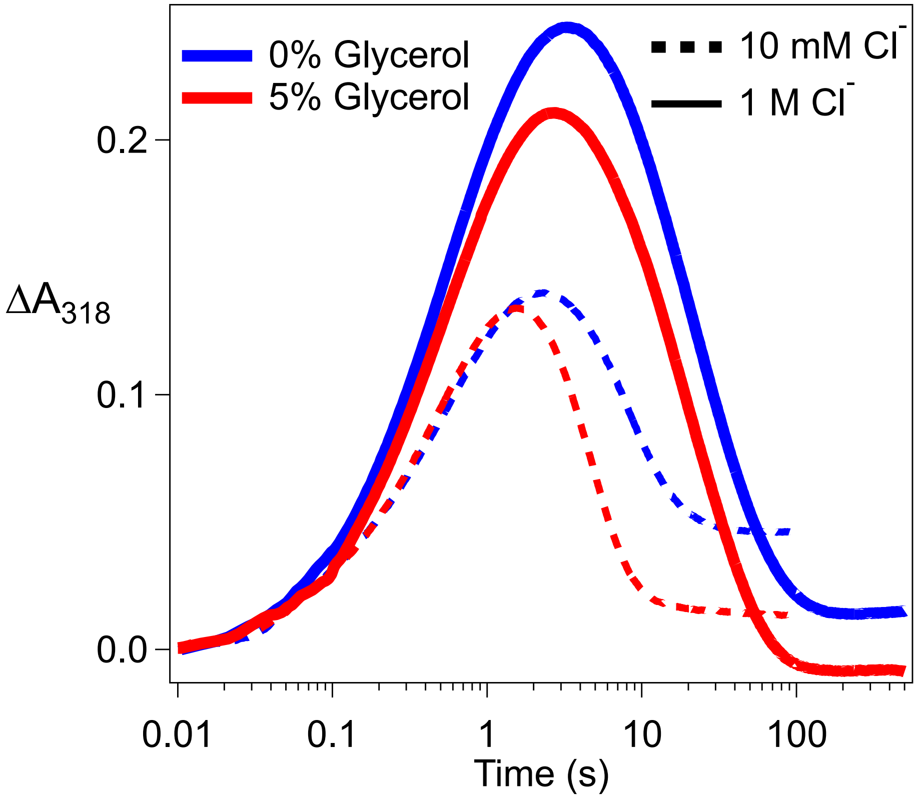


**Figure S12.** Kinetics of the BesD chloroferryl intermediate formation and decay in reactions with $d_4$-L-Lys at 10 mM (*dashed*) and 1 M [$Cl^-$] (*solid*) in the presence (*red*) or absence (*blue*) of glycerol. An anoxic solution of BesD (0.55 mM), $Fe^{II}$ (0.4 mM), 2OG (5 mM), $d_4$-L-Lys (5 mM), and glycerol (0 or 5%, v/v) was mixed at 5°C with an equal volume of air-saturated buffer (~0.18 mM final [$O_2$])

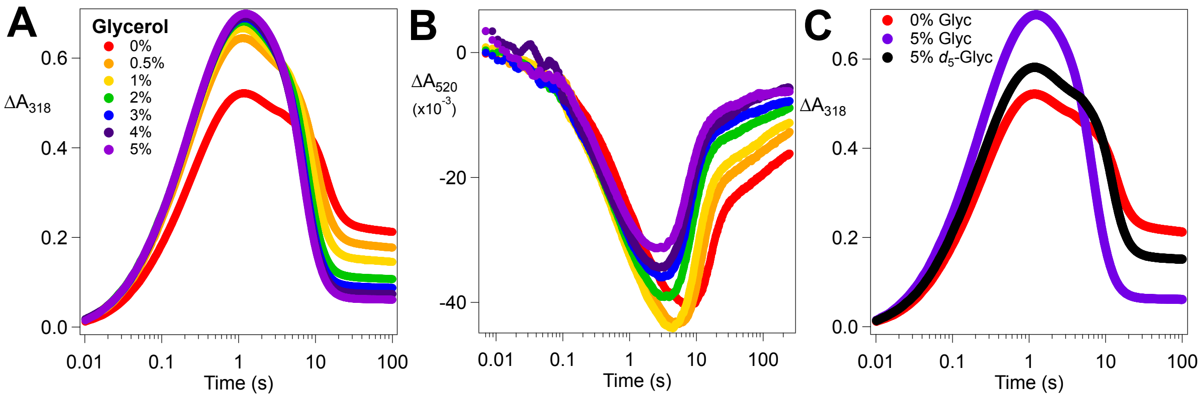


**Figure S13.** Raw kinetic traces of the chloroferryl intermediate formation and decay at (A) 318 nm and (B) 520 nm as a function of glycerol concentration. An anoxic solution of BesD (1.1 mM), $Fe^{II}$ (0.8 mM), 2OG (10 mM), NaCl (100 mM), L-Lys (3 mM), and glycerol (0-5%, v/v) was mixed at 5°C with an equal volume of $O_2$-saturated buffer (~0.9 mM final [$O_2$]). (C) Chloroferryl kinetic traces from (A) for 0% glycerol (*red*) and 5% glycerol (*purple*) compared to a sample containing 5% $d_5$-glycerol (*black*).

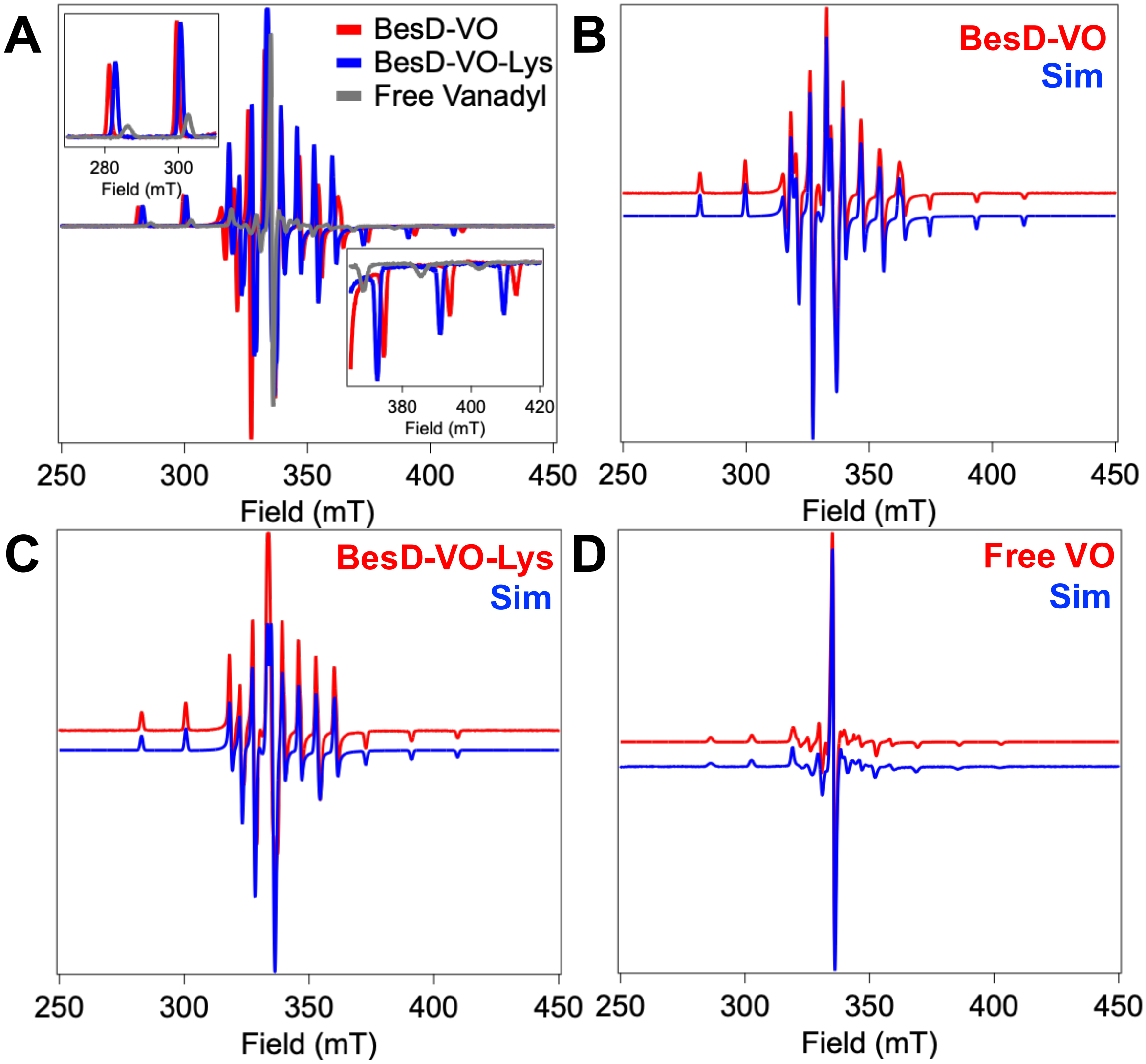


E

| | $g_x$ | $g_y$ | $g_z$ | $A_x$ | $A_y$ | $A_z$ | gStrain-x | gStrain-y | gStrain-z |
|---|---|---|---|---|---|---|---|---|---|
| **BesD-VO** | 1.942 | 1.979 | 1.980 | 511.1 | 190.7 | 177.7 | 0.006 | 0.007 | 0.007 |
| **BesD-VO-Lys** | 1.945 | 1.977 | 1.981 | 492.5 | 165.1 | 164.2 | 0.007 | 0.006 | 0.007 |
| **Free Vanadyl** | 1.960 | 1.979 | 1.994 | 453.8 | 144.2 | 45.1 | 0.012 | 0.009 | 0.005 |

**Figure S14:** Reference spectra and simulations of the three different vanadyl species present in L-Lys and NaCl titration data in Figure S14. (A) Overlay of the three reference spectra with low and high field wings emphasized (*insets*). Experimental and simulated spectrum for (B) BesD-VO, (C) BesD-VO-Lys, and (D) free vanadyl with no protein added to the sample. (E) Simulation parameters for all three species. gStrain is a line shape broadening term used for each g-component.

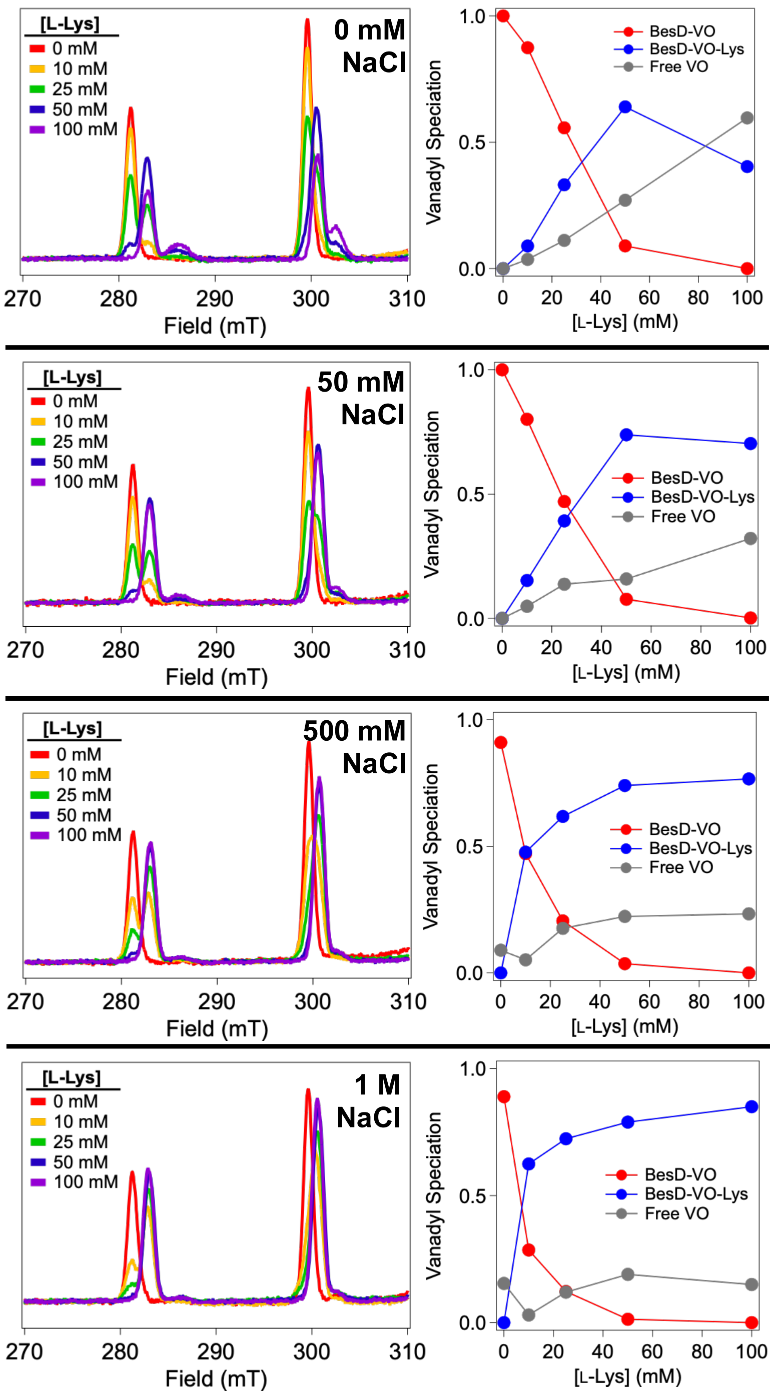


**Figure S15:** EPR spectra of BesD-vanadyl under varying L-Lys and NaCl concentrations. (*Left*) Representative spectra of just the low field section of the vanadyl spectrum are shown for clarity. (*Right*) Population profiles of each vanadyl species present in each sample. Species fractionation was determined using a linear regression fit and reference spectra (Figure S13) for the three species in the software package Igor Pro 9. Sample information and collection parameters is listed in the Material and Methods.

A non-protein-based vanadyl species (Free VO) is observed in the presence of high concentrations of L-Lys and low NaCl likely due to L-Lys chelation in solution which subsequently prevents coordination by BesD.

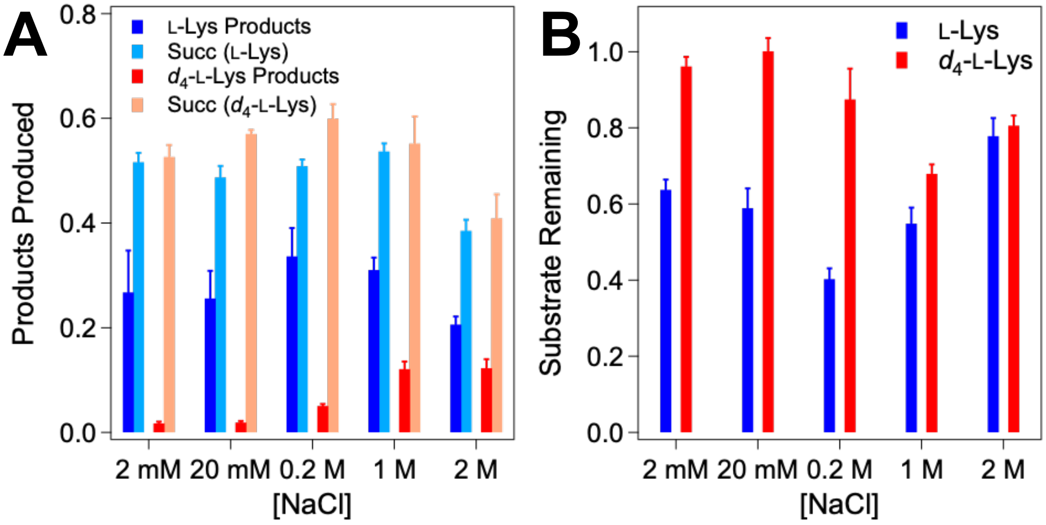


**Figure S16.** (A) Normalized products produced and (B) remaining substrate by multi-turnover assays with L-Lys and $d_4$-L-Lys at varying concentrations of NaCl at 22°C. L-Lys/$d_4$-L-Lys derived product and substrate remaining peak areas were normalized to an internal standard of the opposite isotopologue added post reaction quench. Succinate produced was normalized to an internal standard of $d_4$-succinate added post reaction quench. For reaction conditions, see the Materials and Methods.

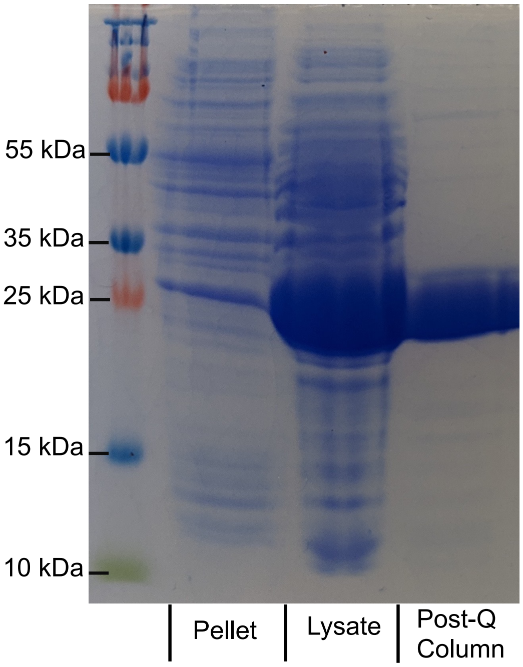


**Figure S17.** Representative SDS-PAGE gel at different purification steps of *Sl* BesD.

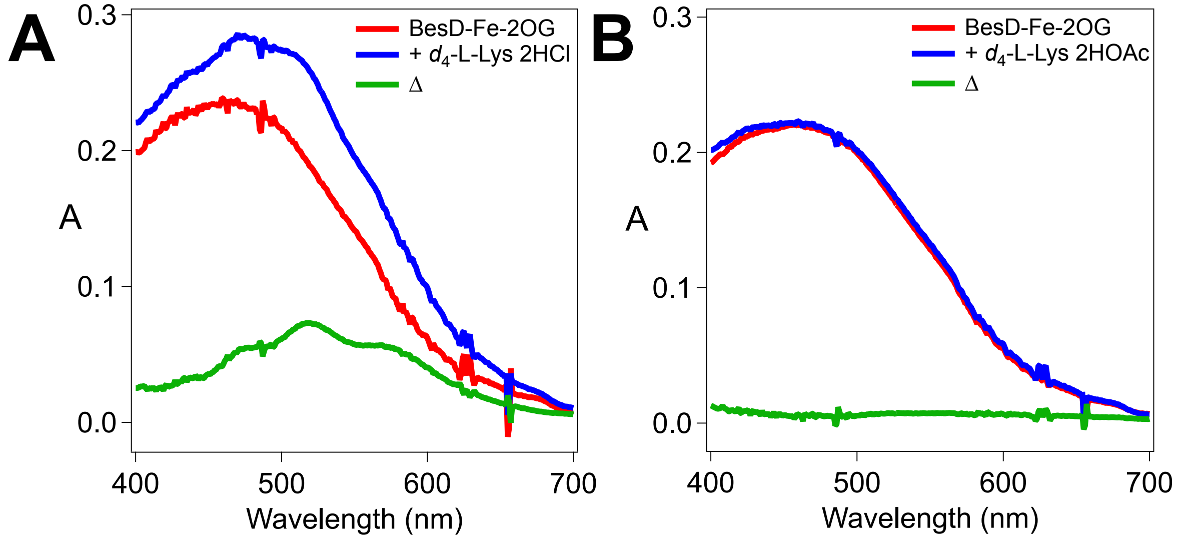


**Figure S18.** (A) UV-Vis absorbance spectra of BesD•$Fe^{II}$•2OG (*red*) before and (*blue*) after addition of $d_4$-L-Lys•2HCl to a final concentration of 30 mM. Difference spectrum is denoted in *green*. (B) UV-Vis absorbance spectra of BesD•$Fe^{II}$•2OG (*red*) before and (*blue*) after addition of $d_4$-L-Lys•2HOAc stock (after ion exchange treatment) to a final concentration of 30 mM. Difference spectrum is denoted in *green*.

**Supplementary References**

(1) DuPont. *Ion Exchange Resins Selectivity*; Tech Fact Form No. 45-D01458-en, Rev. 2; 2019; pp 1–4.
(2) Silakov, A.; Epel, B. Kazan Viewer: Data Processing Software for Matlab, 2022. https://github.com/AlexeySilakov/KazanViewer.
(3) Ravi, N.; Bollinger, J. M.; Huynh, B. H.; Stubbe, J.; Edmondson, D. E. Mechanism of Assembly of the Tyrosyl Radical-Diiron(III) Cofactor of E. Coli Ribonucleotide Reductase: 1. Moessbauer Characterization of the Diferric Radical Precursor. *J. Am. Chem. Soc.* **1994**, *116* (18), 8007–8014. https://doi.org/10.1021/ja00097a007.
(4) Stoll, S.; Schweiger, A. EasySpin, a Comprehensive Software Package for Spectral Simulation and Analysis in EPR. *J. Magn. Reson.* **2006**, *178* (1), 42–55. https://doi.org/10.1016/j.jmr.2005.08.013.
(5) Marchand, J. A.; Neugebauer, M. E.; Ing, M. C.; Lin, C.-I.; Pelton, J. G.; Chang, M. C. Y. Discovery of a Pathway for Terminal-Alkyne Amino Acid Biosynthesis. *Nature* **2019**, *567* (7748), 420–424. https://doi.org/10.1038/s41586-019-1020-y.
